# From sewage to shoreline: Tracing antibiotic resistance gene trends through tropical island wastewater treatment pathways

**DOI:** 10.64898/2026.07.27.740957

**Authors:** Maria Alexa, Aleksandra Kovacevic, Mélanie Pimenta, Degrâce Batantou Mabandza, Thomas U. Berendonk, Sébastien Breurec, Christophe Dagot, Bich-Tram Huynh, Lulla Opatowski

## Abstract

Wastewater is a key reservoir and transmission route for antibiotic resistance genes (ARGs), enabling their spread from influent to effluent and into receiving environments. However, how combined selective pressures (antibiotics, biocides, heavy metals, pharmaceuticals) influence resistant bacteria and ARG persistence over space and time remains poorly understood. Likewise, the role of the wastewater microbiome in ARG dynamics is still unclear, as few studies integrate microbiome shifts with chemical and environmental drivers. Here, we investigated how microbiome dynamics, chemical exposures, and environmental conditions shape clinically relevant ARG dynamics from sewage to receiving environments in Guadeloupe, French Caribbean. We analysed data collected from three wastewater continuums, (hospital-based, domestic, touristic) over four campaigns (September 2021–February 2023). We characterised ARG and microbiome composition spatiotemporal patterns and used a mixed-effect model to investigate ARG associations with potential drivers, including exposome factors, microbiome dissimilarity and environmental factors. Several ARGs were negatively associated with microbiome dissimilarity (Bray–Curtis distances) (*aac(6’)-Ib, aph(3’)-III, bla*_SHV_*, bla*_TEM_*, intI1, qnrS, sul1* and *tetM*). Negative associations were also observed between upstream–downstream differences in anti-inflammatory drug concentrations and the abundance of *aac(6’)-Ib*, *aph(3’)-III*, *bla*_CTX-M_, *ermB*, *intI1*, and *tetM*. In contrast, ARG relative abundance was positively associated with upstream–downstream differences in antibiotic concentrations, suggesting selection along the continuum. These findings indicate that ARG dissemination along wastewater-to-coastal pathways is shaped by opposing processes, with microbiome turnover potentially limiting ARG persistence while chemical gradients promote specific gene enrichment. The outcome is ARG-specific, with implications for antimicrobial resistance risks associated with recreational waters, seafood consumption, and coastal ecosystem interactions.

## Introduction

Antimicrobial resistance (AMR) is one of the most pressing global health challenges of the 21st century, with far-reaching implications for human, animal, and environmental health^1^. A comprehensive global analysis estimated that AMR was directly responsible for 1.27 million deaths and associated with nearly 5 million deaths worldwide in 2019, surpassing the mortality burden of HIV/AIDS and malaria ^2^.

The environment plays a key role in the dissemination of AMR, acting both as a natural reservoir for antibiotic resistant genes (ARGs) and mobile genetic elements (MGEs), and as a pathway for their spread ^3–6^. This is amplified under anthropogenic pressures, particularly through clinical antibiotic use ^7,8^.

Wastewater acts as a major hub for antibiotic resistant bacteria (ARBs), antibiotic resistant genes (ARGs) and chemical pollutants, reflecting anthropogenic activities. By collecting urban wastewater, wastewater treatment plants (WWTPs) integrate a diverse mix of chemical and biological pollutants from domestic, hospital, and (sometimes) stormwater sources before treatment^9^. Although conventional treatment processes reduce partially microbial loads, they are not designed to specifically eliminate ARGs and MGEs^10^. Consequently, tracking the flow of ARGs, MGEs, and associated pollutants from influent to effluent and into receiving environments, a pathway defined here as a continuum, is key to understanding how resistance evolves and spreads within these systems. However, to date, existing studies have largely focused on antibiotics and ARGs along wastewater discharge pathways ^11–14^ while the role of other relevant pollutants remains less explored.

The dynamics of ARGs and MGEs in wastewater can be shaped by the action of selective agents, including antibiotics, biocides, heavy metals, and pharmaceuticals, here defined as the exposome^15^. These contaminants can favour bacteria harbouring ARGs and MGEs through co- and cross-selection mechanisms, thereby enhancing the selection and persistence of antibiotic-resistant bacteria (ARB) in the water ^16–19^. In parallel, the wastewater microbiome potentially plays a fundamental role in shaping ARG persistence and dissemination, with strong associations observed between bacterial community composition and ARG profiles^20^. Emerging evidence suggests that the microbial community plays a dual role in resistance dissemination ^21^. Although high microbial diversity can impede ARG invasion and reduce resistance establishment in stable environments such as soils ^22^, environmental stressors, including heavy metals, and temperature fluctuations, may disrupt this protective effect and instead facilitate ARG persistence and spread ^23^. Despite these insights, few studies have simultaneously examined microbiome dynamics, chemical pollutants, and environmental variables across spatial and temporal gradients to disentangle their combined effects on the resistome.

This knowledge gap is particularly pronounced in wastewater to coastal continuums, where anthropogenic pollution varies widely. For example, hospital effluents carry high levels of ARGs ^24^, whereas downstream environments such as coastal waters and mangroves may exhibit lower or more variable levels. Mangroves, in particular, present a paradox: some studies suggest they act as natural active filters that reduce AMR, while others indicate they may serve as reservoirs of resistance ^25^. Investigating these contrasting observations requires a spatiotemporal framework that captures interactions between the microbiome and exposome across the full environmental continuum. The release of ARGs and ARB into coastal environments raises concerns regarding their transmission through recreational water use, seafood consumption, and ecosystem interactions, reinforcing the need to understand AMR dynamics across the entire wastewater continuum.

In this study, we address these gaps through a comprehensive spatiotemporal analysis of microbiomes, ARGs, and MGEs across three distinct wastewater continuums between September 2021 and February 2023. We compare hospital-based, domestic, and touristic wastewater continuums, in Guadeloupe, a Caribbean island, tracking transitions from raw influent through treatment processes to final discharge into coastal waters and mangroves. We further assess shifts in microbial community composition and changes in exposome factors, including antibiotics, biocides, heavy metals, and anti-inflammatory drugs, to evaluate how these combined drivers influence the persistence, selection, and dissemination of resistance. By using a full wastewater continuum perspective, this work aims to provide a more integrated understanding of resistance dynamics within wastewater systems.

## Results

### Microbiome diversity across wastewater continuums

Along each continuum, microbial communities showed a progressive taxonomic turnover (Figure 1A) from sewage to coastal environments, with the highest degree of community turnover occurring between effluent and ocean or mangrove samples. In alpha diversity analyses (Figure 1B), significant reductions in Chao1 richness were detected only between effluent and ocean in the hospital continuum, from an average of 114 to 58 (p ≤ 0.05, Bonferroni-corrected Wilcoxon test), and between effluent and mangrove in the touristic continuum from an average of 123 to 76 (p ≤ 0.05, Bonferroni-corrected Wilcoxon test) (Supplementary Table S5), suggesting less of a mixture of distinct microbial communities at end points. The Shannon diversity index, which integrates species richness and evenness, also showed significant declines between effluent and ocean samples in the hospital continuum (p ≤ 0.05, Wilcoxon test with Bonferroni correction) (Supplementary Table S6) but unsignificant decline between effluent and mangrove in the touristic continuum.

**Figure 1.**
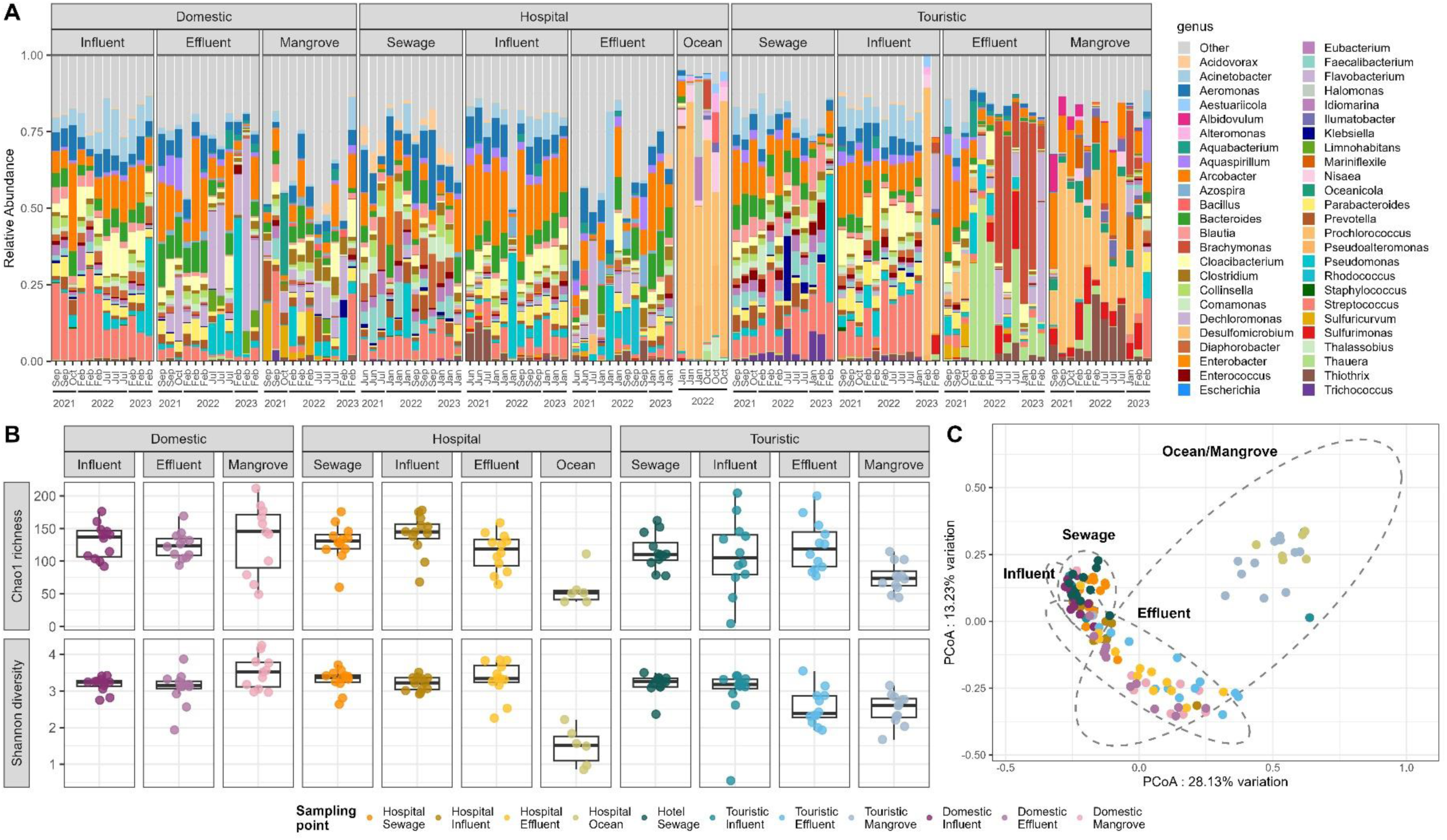
Microbiome structure and diversity across wastewater continuums. **A**. Taxonomic composition of microbial communities across sampling sites and continuums. Top 10 most abundant genera per sampling point, with a MRA >1% are present. ESKAPEE genera are present as well. ‘Other’ captures all other genera. **B**. Alpha diversity of microbial communities across sampling sites (Chao1 richness and Shannon diversity indices across all samples). **C.** Beta diversity of microbial communities based on Bray-Curtis dissimilarity. Principal coordinates analysis (PCoA) showing clustering of samples by microbial community composition. Ellipses only for visual purposes.

Beta diversity analyses further revealed clear spatial structuring, with coastal and wastewater-associated communities separating along the first principal coordinate axis (Figure 1C). This axis was primarily driven by Prochlorococcus (Supplementary Figure S9), a dominant marine cyanobacterium in open ocean waters^26^, which was part of only the hospital ocean and touristic mangrove core communities, with mean relative abundances of 59.9% and 28.0%, respectively. Sampling location explained 44% of the variance in microbial community composition (adonis2, p ≤ 0.001).

Across all samples, 616 bacterial genera were detected. Core genera were shared among the three continuums (Supplementary Table S4), indicating a common wastewater-associated microbiome. At the genus level, *Arcobacter* was detected in the core community of all sampling points except the hospital ocean samples, with mean relative abundances ranging from 20.1% in the hospital influent to 5.7% in the domestic mangrove. *Streptococcus* was part of the core communities of all sewage and influent samples, with mean relative abundances of 15.9% in the domestic influent and 5.2% in the hospital influent. *Aeromonas*, an opportunistic pathogen that can act as a reservoir of last-line antibiotic resistance genes ^27^ was also present, with the highest mean relative abundance in hospital sewage (10.5%). *Bacteroides*, a major member of the gut microbiota, was shared across sewage, influent and effluent samples, except for the touristic effluent*. Pseudomonas* and *Acinetobacter* were also abundant, with the highest combined mean relative abundance in the domestic influent (8.7%), and both included members of the ESKAPEE group, such as *Acinetobacter baumannii* and *Pseudomonas aeruginosa* ^28^.

Despite this shared core, each continuum also showed distinct local core communities. Hospital sewage had a specific core microbiome characterised by *Diaphorobacter* (7.9%), *Faecalibacterium* (5.2%), and *Eubacterium* (3.7%), reflecting its source-specific profile. The touristic effluent showed marked temporal variation across sampling campaigns: early periods were dominated by *Thauera* (mean relative abundance 15.5%), whereas later campaigns showed increased abundance of *Brachymonas* (18.7%), suggesting a possible temporal restructuring of the effluent microbiome.

### Spatiotemporal trends of antimicrobial resistance genes across wastewater continuums

Trends observed along the hospital continuum reveal that hospital sewage and hospital influent (a mixture of hospital sewage and domestic wastewater) consistently had the highest relative abundance across nearly all clinically relevant genes compared to the domestic and touristic continuums. Across all three wastewater continuums, influent and upstream sources (hospital sewage or touristic sewage) showed the highest relative abundances of ARGs, particularly for mobile and clinically relevant genes such as *ermB*, *intI1*, *tetM*, and *sul1* (Figure 2). Wastewater treatment resulted in modest reductions.

**Figure 2.**
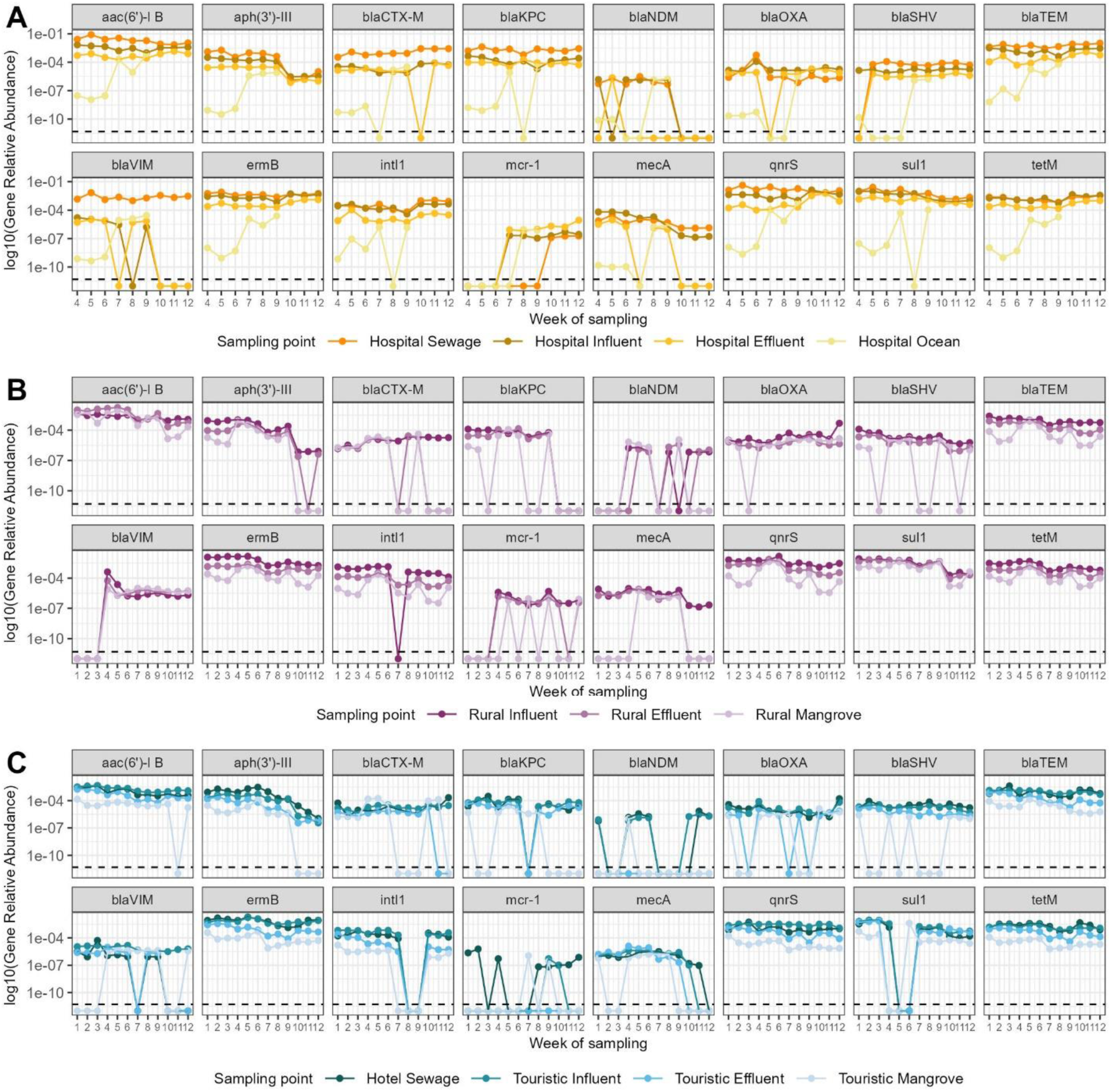
Spatiotemporal distribution and persistence of clinically relevant antibiotic resistance genes across wastewater continuums in Guadeloupe. **A.** Hospital wastewater continuum **B.** Domestic wastewater continuum **C.** Touristic wastewater continuum. Data represent log₁₀-transformed relative abundances of 16 clinically relevant antibiotic resistance genes. The dashed line represents the limit of quantification. Colours indicate the location point. The x-axis represents the log₁₀-transformed relative abundance and y-axis the week of sampling. Week 1 to 3 represent the first sampling campaign, 4 to 6 the second sampling campaign, 7 to 9 the third sampling campaign and 10 to 12 the fourth sampling campaign.

In the receiving environments, ocean and mangroves, the ARGs *bla*_TEM_, *ermB*, *qnrS*, and *tetM* showed a 100% detection frequency, being detected above the limit of detection in all samples collected. Additional ARGs were detected at 100% detection frequency only in domestic mangrove and hospital ocean, including *aac(6’)-Ib* and *sul1*. It is worth noting that the mean total relative abundance was lower in receiving environments compared with upstream wastewater samples (7.2e-4 in ocean compared with 2.0e-1 in hospital sewage; 2.4e-2 in domestic mangrove compared with 1.0e-1 in domestic influent; and 4.2e-3 in touristic mangrove compared with 9.6e-2 in hotel sewage), potentially due to dilution.

### Clustering of antimicrobial resistance genes

Analysis of Bray-Curtis dissimilarities in resistome composition across all ARGs revealed geographical structuring by sample location, with sampling point accounting for 43% of the observed variation (adonis2, p < 0.001). Samples from the touristic mangrove and hospital ocean sites were distinctly separated from other samples along the first principal coordinate (Figure 3A).

**Figure 3.**
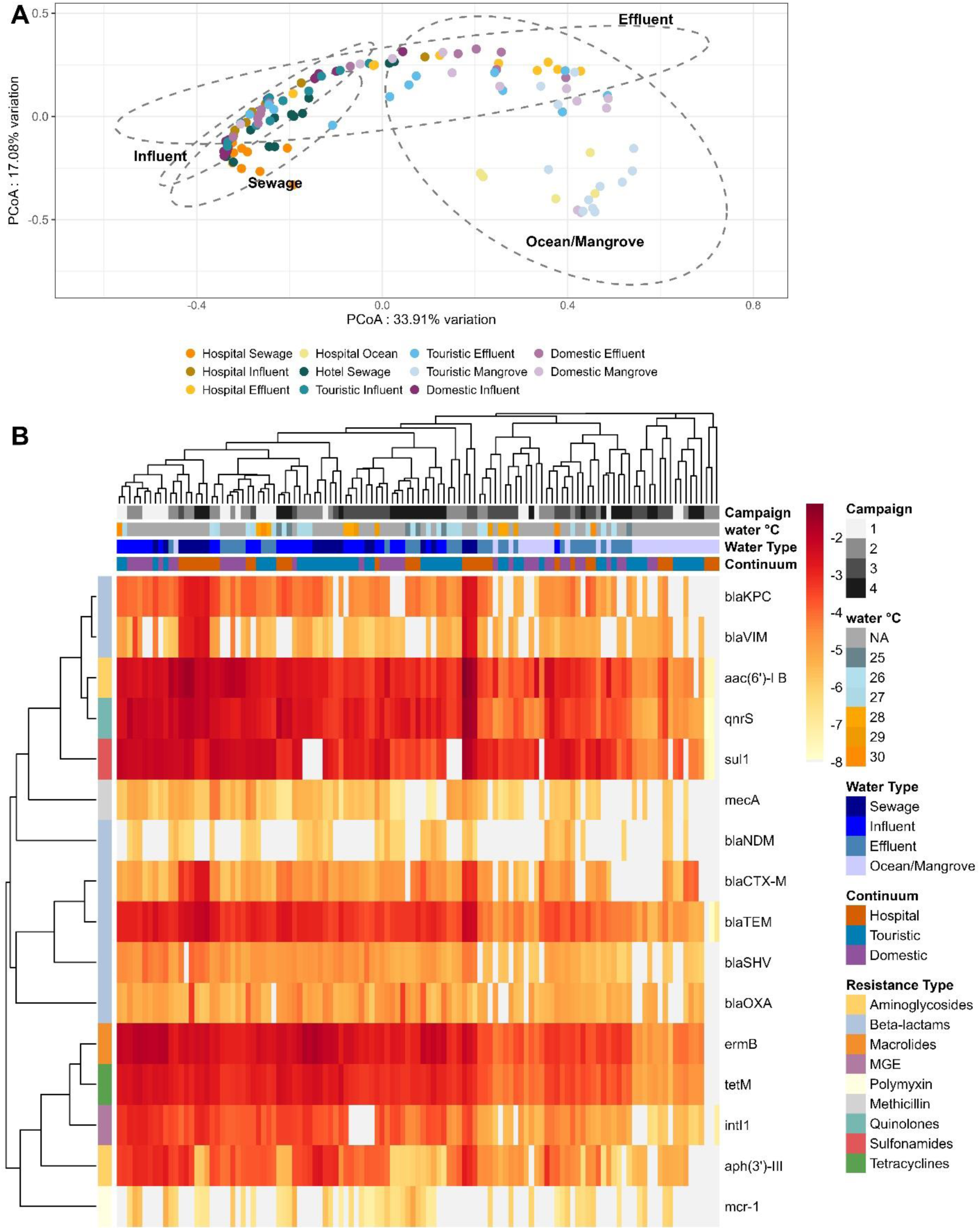
Resistome clustering of genes of interest. **A**. Principal coordinate analysis (PCoA) using the resistome Bray-Curtis dissimilarity matrix. **B**. Clustered heatmap of 16 clinically relevant genes of interest. Colors in the heatmap represent *log*_10_ transformed relative abundance. Columns represent each sample collected and are clustered based on the full resistome Bray-Curtis dissimilarity matrix. Rows represent each gene of clinical interest and are clustered based on Pearson correlation using complete-linkage. Annotation colors for columns (samples) are based on water temperature, water type (sewage, influent, effluent and ocean/mangrove), continuum and sampling campaign. Annotation colors for row (genes) are based on the resistance type.

Using heatmaps, we found that this separation was primarily driven by higher relative abundances of genes such as *aac(6’)-Ib*, *qnrS, sul1*, *bla*_TEM_, *ermB*, *tetM,* and MGE *intI1*, which were consistently prevalent in sewage and influent samples across all continuums (Figure 3B). Genes *bla*_KPC_*, bla*_VIM_*, bla*_CTX-M_ had their highest relative abundance in hospital sewage. In contrast, touristic mangrove and ocean samples showed lower overall ARGs abundances, with many genes falling below detection limits. Overall, clustering patterns primarily reflected sample type, distinguishing sewage, influent, effluent, and ocean/mangrove samples.

When considering clustering across all resistance genes (Supplementary Figure S11), *ermB* and *tetM* were consistently grouped together. Additional clusters of clinically relevant genes were observed, including *bla*_VIM_ with *bla*_KPC_, and *qnrS* with *aac(6’)-Ib*. Additionally, *sul1* clustered together with *IS6* group and *intI1* with *ISEc9*.

### Environmental and microbiome drivers of ARG abundance

A mixed-effects model was used to analyse the spatiotemporal dynamics of each clinically relevant ARG independently. Univariate analyses results and mixed-effect model results are summarized in Tables S9 and S10 (Supplementary Materials) and shown in Figure 4.

**Figure 4.**
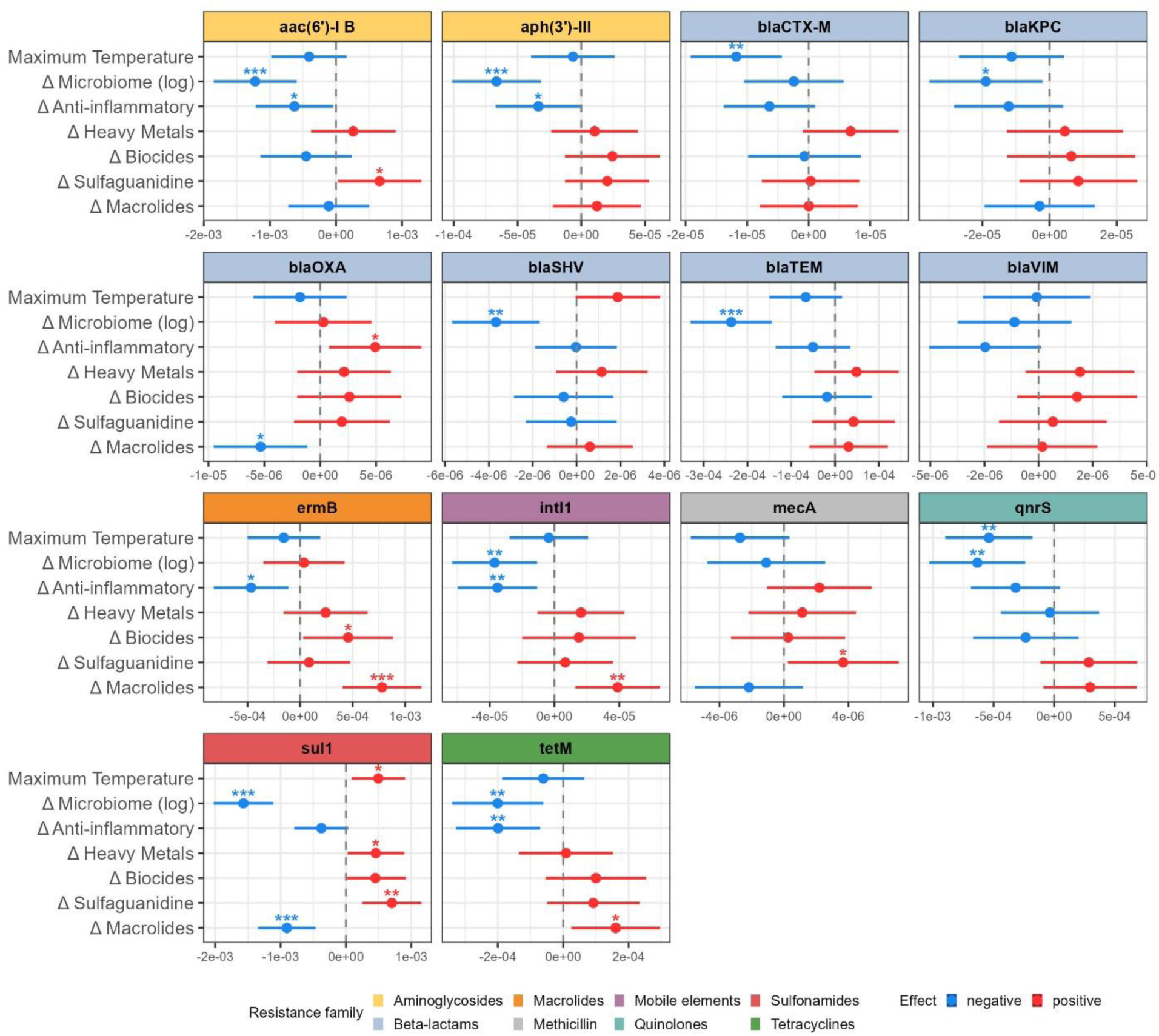
Estimated effects of covariates on the abundance of antibiotic resistance genes (ARGs) from gene-specific mixed-effects models. Points show estimates of regression coefficients from 14 mixed-effects models (one per ARG of interest), with horizontal bars representing coefficients 95% credible intervals. The dashed vertical line indicates no effect (*β* = 0). Blue and red denote negative and positive associations, respectively. Statistical significance is indicated by asterisks (*p < 0.05; **p < 0.01; ***p < 0.001). ARGs are grouped by the resistance family. Δ*Microbiome* corresponds to the Bray–Curtis dissimilarity between upstream and downstream sites. Other covariates are expressed as downstream minus upstream values. The carbapenemase gene *bla_NDM_* and the colistin resistance gene *mcr-1* were excluded from the analyses due to insufficient detection frequency.

First, microbiome dissimilarity (Bray–Curtis) was negatively associated with the relative abundance of 12 out of 14 clinically relevant ARGs. This association was statistically significant for eight ARGs (*aac(6’)-Ib*, *aph(3’)-III*, *bla*_KPC_, *bla*_SHV_, *bla*_TEM_, *intI1*, *qnrS*, *sul1,* and *tetM*) (Figure 4) and not significant for *bla*_CTX-M_, *bla*_VIM_, and *mecA*. Second, upstream–downstream contrasts in anti-inflammatory drug concentrations were significantly negatively associated with *aac(6’)-Ib*, *aph(3’)-III, ermB, intI1*and *tetM.* Third, maximum temperature was negatively associated with *bla*_CTX-M_ and *qnrS* as well as positively associated with *sul1*. Moreover, upstream–downstream differences in antibiotic concentrations along the continuum were also positively associated with ARG abundance. Differences in macrolide concentrations were positively and significantly associated with *ermB, intI1* and *tetM,* but negatively associated with *bla*_OXA_ and *sul1*. Specifically, the sulfaguanidine gradients were positively associated with all ARGs, except *bla*_SHV_, although this was statistically significant only for *aac(6’)-Ib*, *mecA*, and *sul1*. Finally, differences in heavy metal and biocide concentrations were generally positively associated with ARG abundance, although these associations were not statistically significant, except for the association between biocides and *ermB* and heavy metals and *sul1*.

For most ARGs of interest, except *bla*_VIM_, *bla*_KPC_, and *mecA*, the multivariable model (M-multi) had lower AIC values than the null model (M-null) (Supplementary Table S11), suggesting that the multivariable model was better at explaining the data.

For ESKAPEE genera, only analyses of *Enterococcus* and *Staphylococcus* showed lower AIC values with the multivariable models compared to the null model (Supplementary Table S12) suggesting that the included covariates improve the model fit of these two genera, unlike for the other ESKAPEE taxa where the covariates did not add explanatory power beyond the null model. Microbiome dissimilarity (Bray–Curtis) was significantly negatively associated with *Staphylococcus* relative abundance (Supplementary Figure S15).

## Discussion

Using longitudinal data from three wastewater continuums (hospital, domestic and touristic) up to their coastal receiving end points, we analysed the spatial and temporal dynamics of ARGs and their potential determinants in Guadeloupe, including the wastewater exposome (antibiotics, anti-inflammatories, biocides and heavy metals) and the microbiome. Our results showed that the influence of these determinants was strongly ARG-specific, with contrasting factors shaping the fate of individual resistance markers. While chemical exposures, including antibiotic, biocide, and metal gradients, may provide selective pressures favouring the persistence of certain ARGs, microbiome turnover appeared to act as an ecological barrier limiting the maintenance of others. Consequently, no single driver universally explained ARG dynamics across wastewater continuums; rather, the relative contribution of selective pressures and ecological constraints varied among ARGs.

Spatial temporal dynamics observed in the hospital wastewater continuum reveal that raw hospital sewage and hospital influent consistently showed the highest ARG and MGE levels across nearly all genes of interest, compared to the domestic and touristic continuums. For all three continuums, effluent levels were typically only slightly lower. This could indicate that there is only a partial treatment success in removing resistant genes and suggest constant input of ARGs and MGEs from both clinical sources and domestic sources. The consistent detection of *bla*_TEM_, *ermB*, *qnrS*, and *tetM* across all receiving environments highlights their potential value as targets for monitoring temporal and spatial variations in the ocean and mangrove resistomes. Similar ARGs have frequently been reported in recreational and coastal waters, reflecting their widespread distribution and association with anthropogenic inputs^42^. However, relative abundance should be interpreted cautiously^43^ in ocean and mangrove samples, where the total relative abundance was low (Supplementary Figure S4).

Our estimates indicated that changes in microbiome composition along the continuums were significantly and negatively associated with the relative abundance of eight ARGs (*aac(6’)-Ib, aph(3’)-III, bla*_SHV_*, bla*_TEM_*, intI1, qnrS, sul1 and tetM*). These findings suggest that community turnover, and, by extension, increased ecological diversity, may act as an ecological barrier to ARG persistence. In our case, higher dissimilarity between upstream and downstream communities may reflect enhanced ecological competition or niche restructuring, thereby limiting the establishment and persistence of pathogens carrying clinically relevant ARGs. In concordance with previous studies in soils, Klümper et al. (2024)^22^ reported that higher diversity, evenness and richness were significantly negatively correlated with the relative abundance of more than 85% of ARGs, and proposed that microbial diversity can reduce the persistence time of immigrating ARGs. Although their work focused on soils rather than aquatic systems, our results suggest that similar ecological mechanisms may operate in wastewater continuums. Our findings also suggest that the microbiome potentially plays a central role in ARG dynamics along these continuums and should be considered alongside chemical exposures in risk assessment and management strategies.

One important finding of our study was that increases in antibiotic concentrations between upstream and downstream sites were generally positively associated with the downstream relative abundance of clinically relevant ARGs. In other words, sites with larger increases in antibiotic concentrations tended to exhibit higher downstream ARG relative abundances. Specifically, increases in sulfaguanidine concentrations were positively associated with all ARGs, with statistically significant associations for *aac(6’)-Ib, mecA* and *sul1*. The positive association between sulfaguanidine and *sul1* is consistent with previous studies reporting links between sulfonamides and *sul1* in surface waters and WWTP effluents ^29,30^. In concordance with these studies, our results support the idea that exposure to antibiotics, even at relatively low concentrations, can select for ARGs over space and time. Increases in macrolide concentrations between upstream and downstream sites were also positively and significantly associated with *ermB*, *intI1* and *tetM*. For *intI1*, this is consistent with Bijlsma et al. (2024)^31^, who reported a comparable positive association with macrolides and interpreted it as co-selection. In contrast, the positive association we observed for *ermB* was not found in that study, which may reflect differences in exposure patterns, co-occurring contaminants, or local microbial community structure.

Beyond these individual associations, *ermB* and *tetM* clustered together in our dataset (Supp. Fig. S10), pointing to a shared dissemination route rather than independent selection. (Fig. 5). This aligns with previous findings in Spanish wastewater, where *tetM* was positively correlated with macrolides^31^. The co-occurrence of *ermB* and *tetM* is likely driven by their frequent co-localization on mobile genetic elements, particularly conjugative transposons, as described for *Streptococcus pyogenes* ^32^. Given that *Streptococcus* was among the 15 most abundant genera in our dataset (Fig. 2A), it may contribute to the co-occurrence and co-selection of these genes along the continuums. These findings suggest that selection by macrolides may act not only on individual ARGs, but also on ARG assemblages carried on shared mobile genetic elements.

**Figure 5.**
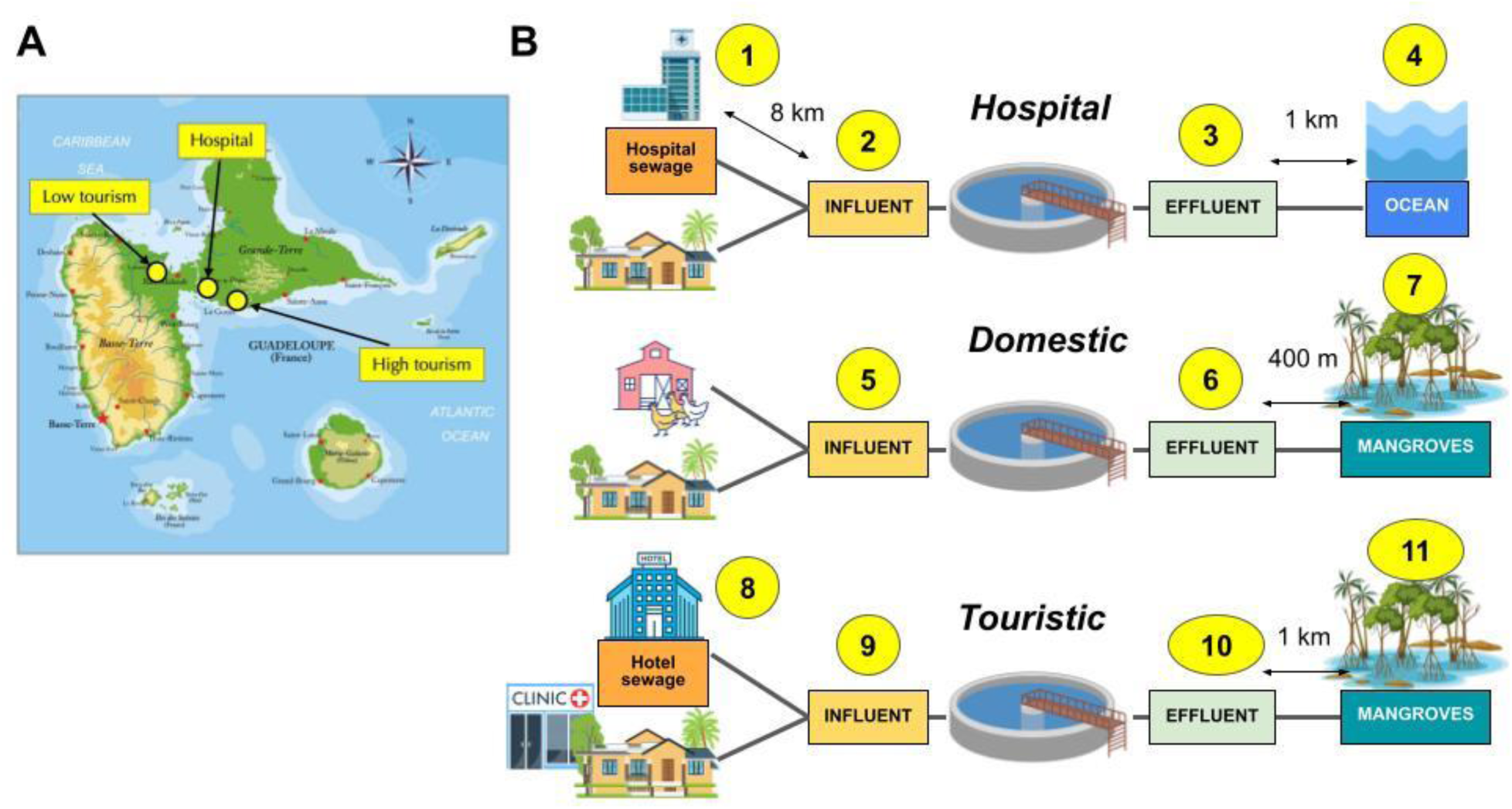
Study location and sampling design. **A.** Map of Guadeloupe showing the locations of the three wastewater continuums: hospital (Pointe-à-Pitre), domestic (Lamentin), and touristic (Le Gosier), with associated sampling campaign periods (C1 – C4). **B.** Schematic representation of sampling points along the hospital, domestic, and touristic wastewater continuums, showing the flow from sources through treatment and into receiving environments (ocean or mangroves). For more details, see Table S1, Supplementary Material.

We found that maximum air temperature was positively associated *sul1* and negatively associated with the relative abundance of *bla*_CTX-M_ and *qnrS*. The negative association contrasts with some previously reported findings^33,34^. It may be explained by the fact that maximum temperature was not measured at each sample site but only at the three WWTPs. To include this variable as an explanatory factor in the mixed-effects model, we assumed that within each continuum and sampling week, maximum temperature was homogeneous across all sampling points. Given the relatively short distances between sampling points within each continuum (up to 3 km), this assumption is likely. However, increasing temperature has been reported to have a dual impact, both being associated with increased ARG abundance in naturally occurring biofilms and with reduced establishment of newly invading ARGs ^35^.

Our results showed that changes in anti-inflammatory drug concentrations were significantly negatively associated with several ARGs *(aac(6’)-Ib, aph(3’)-III, bla*_CTX-M_*, ermB, intI1 and tetM)*. This pattern contrasts with recent experimental work by Zhang et al. (2024)^36^, who reported that ketoprofen can accelerate the environmental spread of ARGs by increasing cell-to-cell contact and inducing an SOS response. However, this discrepancy is likely explained by the markedly different exposure levels used in the two contexts. In our study, anti-inflammatory drugs were detected at environmentally relevant concentrations in the µg/L range, with maximum values of 1.8 µg/L for ketoprofen (hospital sewage; Supplementary Figure S8), 3.3 µg/L for diclofenac, and 5.2 µg/L for ibuprofen. In contrast, Zhang et al. (2024)^36^ observed enhanced conjugative transfer at much higher concentrations, with peak effects at 10 mg/L ketoprofen and the highest transfer frequency at 50 mg/L, which is several orders of magnitude above the concentrations measured in our samples. Further studies are needed to determine the concentration thresholds at which different anti-inflammatory drugs can promote ARG dissemination in complex environmental settings, including wastewater and mangroves.

Finally, changes in heavy metal and biocide concentrations were generally positively associated with ARG abundance, although most of these associations did not reach statistical significance. The main exception was a significant positive association between biocides and *ermB*. This result is consistent with the concept of co-selection, whereby heavy metals and biocides select for mobile genetic elements carrying both resistance genes to these compounds and ARGs ^16,17,19,37^. Our estimates, although often non-significant, point towards a broad pattern of positive relationships between these co-stressors and ARGs, and they highlight the need to consider cumulative and interacting selection pressures rather than antibiotics alone ^38^.

In parallel with these chemical drivers, we observed microbiome marked spatial and temporal shifts along the continuums, particularly in the touristic effluent. The observed temporal shift in the tourist continuum, with an early predominance of *Thauera* followed by an increased relative abundance of *Brachymonas*, suggests possible reorganization of the microbial community over time. *Thauera* belongs to the Rhodocyclaceae family, which is a core bacterial group in activated sludge globally, as shown in a survey of 269 WWTPs across six continents ^39^. *Thauera* has also been reported to be highly competitive in low-carbon wastewater treatment ^40^ supporting its dominance under certain operational or environmental conditions. Other studies have observed seasonal shifts of microbiome communities in WWTPs and have proposed community stability as a potential control parameter for WWTP maintenance ^41^, highlighting the need for longer time series that integrate operational and environmental metadata to better resolve the drivers of these microbiome temporal shifts.

Several aspects of the study design should be considered when interpreting our findings, while also providing useful directions for future research. First, environmental variables such as maximum temperature were measured at the WWTP level rather than at each individual sampling point. Because sampling points within each continuum were geographically close, environmental conditions were expected to be broadly comparable. Nevertheless, site-specific measurements of maximum temperature, water temperature and other physicochemical variables would provide a more precise characterization of local conditions.

Because compounds within the same chemical class were strongly correlated, we used aggregated class-level measures for antibiotics, biocides, heavy metals, and anti-inflammatory drugs. This approach reduced multicollinearity and enabled the identification of broader class-level patterns, although it may have masked compound-specific effects. Future studies combining larger datasets with compound-specific modelling could help distinguish the contributions of particular contaminants. Additional hydrological and environmental variables, such as water flow, could also be included in future studies, to avoid potential confounding effects and improve understanding of ARG transport and persistence.

Secondly, our choice of mathematical function to describe the relationship between ARG relative abundance and chemical compound concentrations in the mixed-effect model can be discussed. Our primary model related ARG relative abundance to upstream-downstream changes in compound concentration (*RA ∼ δC*). To assess robustness of this approach, we also tested changes in ARG relative abundance as a function of changes in compound concentrations (*δRA ∼ δC*), which yielded weaker and more limited associations, with statistically significant relationships retained only for macrolides–*sul1* and sulfaguanidine–*mecA* combinations (Supplementary Fig. S15).

A further limitation is the absence of detailed information on the bacterial hosts carrying the detected ARGs and MGEs. We analysed genes independently, although some may be co-located on the same bacterial genomes or mobile genetic elements. However, the absence of bacterial host data prevented the investigation of such linkages. This limitation may partly explain why similar compounds were associated with the different ARGs.

Moreover, ESKAPEE-related results should be interpreted with caution, as 16S rRNA gene sequencing provided genus-level taxonomic resolution. Consequently, we could not distinguish pathogenic from non-pathogenic species within the same genera. Future studies should aim for species-level taxonomic resolution.

Finally, our sampling design comprised of four sampling campaigns with three weekly samples per campaign, may have limited statistical power for our analysis. This designed aimed at providing valuable spatial and temporal coverage across the three continuums, but the short duration of each sampling campaign may have limited our ability to capture short-term fluctuations and episodic discharge events. Future studies should consider extending the duration of each sampling campaign, while maintaining similar seasonal coverage, to better characterize temporal variability and improve statistical power.

Despite these limitations, the study provides an integrated assessment of ARG dynamics from wastewater influent through treatment and into receiving coastal environments in three wastewater continuums. By combining, spatiotemporal measurements of ARG relative abundance, microbiome composition, chemical contaminants, and environmental conditions, we provide valuable insights into the associations between ARGs and microbiome dissimilarity, antibiotics, anti-inflammatory drugs and maximum air temperature. Our findings emphasize the importance of considering both ecological processes, including microbial diversity and community turnover, and chemical processes, including exposure gradients and co-selection when assessing the environmental dissemination of clinically relevant ARGs.

Future longitudinal studies incorporating higher-frequency sampling, site-specific environmental measurements and hydrological variables could be integrated into our model to better understand the dynamics of antibiotic resistance in coastal environments. Further research should also assess the extent to which ARGs and antibiotic-resistant bacteria in coastal environments may pose risks for human exposure through recreational water use, seafood consumption, and ecosystem interactions, thereby strengthening links between environmental monitoring and public health.

## Materials and Methods

### Sampling sites and sample collection

Sampling was conducted in Guadeloupe between September 2021 and February 2023. Guadeloupe has a resident population of 384,315 inhabitants (2021) and receives approximately 960,000 visitors annually (2022). To assess the impact of anthropogenic activities on wastewater composition, sampling was conducted across three distinct wastewater continuums, each representing a complete pathway from source to environmental endpoint.

The hospital continuum traced wastewater from its origin at the University Hospital Center of Guadeloupe, starting with hospital sewage, through the mixed wastewater influent (a combination of hospital sewage and domestic wastewater), to the post-treatment effluent and its ultimate discharge into the ocean. The domestic continuum followed wastewater from a small town in a rural area with agricultural activity (including poultry farms), through influent and treatment plant effluent before its final release into a mangrove ecosystem. The touristic continuum captured wastewater from a high-tourism area, starting from hotel sewage and progressing through the wastewater treatment plant influent (a mixture of hotel sewage and other wastewater inputs, notably from a dialysis-specialized clinic), to treatment plant effluent and its ultimate discharge into a mangrove system (Figure 5).

To account for seasonal variability, wastewater samples were collected across four successive sampling campaigns, covering two dry and two rainy seasons. Each campaign consisted of weekly sampling for three consecutive weeks at each point along each continuum. A total of 132 wastewater samples (2L each) were collected over the 12-week sampling period. Detailed descriptions of the sampling sites are provided in Table S1 (Supplementary Materials).

### Sample processing and analysis

Water samples were transported to the laboratory at 4 °C and filtered within 24 h through 0.22 µm membranes (Millipore; Dominique Dutscher, Paris, France). The volume processed depended on sample turbidity and was recorded for every sample. Saturated filters were frozen (– 80 °C) in empty haemolysis tubes and sent on dry ice to Limoges (France), where DNA was extracted with the DNeasy PowerWater kit (Qiagen) following the manufacturer’s protocol. DNA yields were determined with a Qubit dsDNA Broad-Range assay (Invitrogen), adjusted to 10 ng µL⁻¹ and stored at –20 °C until use.

High-throughput qPCR (96.96 BioMark Dynamic Array, Fluidigm) was applied to quantify 80 antibiotic-resistance genes (ARGs; 16 classes), genes conferring resistance to quaternary ammonium compounds (QACs), nine mobile genetic elements (MGEs) and the integron integrase genes *intI1*, *intI2* and *intI3*^44,45^. Universal 16S rRNA primers served as the internal reference. Reactions were run in triplicate; a wastewater sample containing all targets and nuclease-free water were used as positive and negative controls, respectively. The relative abundance was calculated using the 16S rRNA encoding gene with the 2^−ΔCT approach (ΔCT = CT_ARG_ − CT_A6S rRNA_)^44,45^. A quality control filter step was initially applied to assess qPCR replicate variability, as detailed in the TextS1, Figure S1 and Figure S2 (Supplementary Materials). This step was designed to assess variability both within and between filters.

Nine antibiotics, eight biocides and three non-steroidal anti-inflammatory drugs were extracted by solid-phase extraction (Oasis HLB 3 cc, 60 mg; Waters) and quantified in order to get their concentrations by LC–MS/MS (5500 QTRAP, Sciex) operated in multiple reaction monitoring (MRM) mode, as described previously ^46^. Heavy-metal concentrations were obtained with an ICP-MS NexIon 350D (Perkin Elmer) using Sc, Y, In and Re as internal standards. Temperature, precipitation and other climate variables were recorded during each sampling campaign at the location of the three wastewater treatment plants only. Detailed listing of the collected data is provided in Table S2 (Supplementary materials).

This study focuses on 16 clinically important targets: *aac(6′)-Ib, aph(3′)-III, bla*_CTX-M_*, bla*_KPC_*, bla*_NDM_*, bla*_OXA_*, bla*_SHV_*, bla*_TEM_*, bla*_VIM_*, ermB, mecA, mcr-1, qnrS, sul1, tetM* and *intI1*. This study also focuses on ESKAPEE pathogens (*Enterococcus faecium*, *Staphylococcus aureus*, *Klebsiella pneumoniae*, *Acinetobacter baumannii*, *Pseudomonas aeruginosa*, *Enterobacter* spp., and *Escherichia coli*) ^28,47^ at the genus level.

### Statistical analysis

#### Alpha diversity and definition of the bacterial core microbiome

Alpha diversity based on the microbiome dataset was estimated using the Chao1 richness and Shannon diversity indices implemented with the *vegan* package in R.

To identify the bacterial core community at the genus level from the microbiome dataset, genera were classified according to their relative abundance and occurrence patterns (presence/absence) at each sampling point across time. Overall abundant genera were defined based on their mean relative abundance (MRA) ^48^, with a MRA greater than 1% across all samples from a same sampling point ^49^. Ubiquitous genera were defined as those detected in more than 80% of samples ^50^ within a sampling point. Frequently abundant genera were identified based on their relative contribution within each sample: genera were considered abundant in a given sample when they ranked among the dominant taxa, collectively accounting for the top 80% of total relative abundance within that sample ^51^. A genus was then classified as frequently abundant when it met this criterion in at least half of the temporal samples within a sampling point. Those three criteria allowed us to define a core community for each sampling point, as done in previous studies^20^.

#### Beta diversity and ordination of microbiome and resistome

Beta diversity was assessed by computing Bray–Curtis dissimilarity matrices for both the microbiome and resistome datasets using the *vegdist* function.

Ordination analysis was performed using principal coordinates analysis (PCoA) with the *wcmdscale* function to visualize patterns of both microbiome and resistome dissimilarity among samples. The effect of sampling location was tested using permutational multivariate analysis of variance (PERMANOVA) implemented with the *adonis2* function.

#### ARG dynamics

To explore spatial patterns in the resistome, relative abundances were log-transformed and visualized in a heatmap using the *pheatmap* package, highlighting genes of clinical interest. Gene-wise clustering was performed using Pearson correlation coefficients, whereas sample clustering was based on Bray–Curtis dissimilarity calculated across all genes. Both gene and sample dendrograms were generated using complete linkage clustering.

#### Spatio temporal dynamics of ARGs

A mixed-effects model was used to analyse ARG relative abundance along the continuums over time and to identify associated factors while accounting for the dependence among sampling locations. The response variable was the relative abundance of ARGs of clinical interest at each downstream sampling point and sampling date. Covariate data included: (i) overall changes in exposome factors between upstream and downstream locations, including biocides, heavy metals, anti-inflammatory drugs, and antibiotics; (ii) Bray–Curtis dissimilarity in microbiome composition between upstream and downstream sites within the same continuum segment and (iii) maximum temperature. The model included fixed effects derived from these covariates and a spatial random effect.

The model was fitted independently for each ARG of clinical interest It is specified as follows:

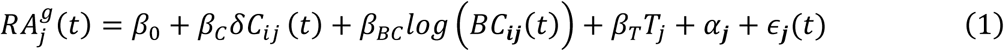

with *α*_*j*_ ∼*N*(*0*, *σ*_*α*_^2^) and *υ*_*j*_(*t*)∼*N*(*0*, *σ*_*υ*_^2^)

Here, *j* is the downstream sampling point, *i* is the corresponding upstream sampling point, and *t* the sampling week. *RA_j_*^*g*^(*t*) represents the relative abundance of a given gene *g* at downstream sampling point *j* during week *t*. *β*_0_is the intercept and the vector *β*_*C*_ contains the fixed-effect parameters of length *p*, where *p* is the number of covariates. The term *δC*_*ij*_ (*t*) denotes the corresponding fixed-effect design matrix (of dimension SxN×p, where S is the total number of downstream sampling points, N is the total number of sampling weeks), representing differences in exposome factors between upstream (*i)* and downstream (*j)* sampling points. *β*_*BC*_ denotes the fixed-effect coefficient associated with Bray–Curtis dissimilarity, and *log*(*BC*_*ij*_(*t*)) the corresponding predictor (dimension SxN×1). *β*_*T*_ denotes the fixed-effect coefficient associated with the maximum temperature, and *T*_*j*_ the corresponding predictor (dimension SxN×1). The term *α*_*j*_ represents the sampling point–specific random intercept, assumed to follow a normal distribution. The residual error term is denoted by *ε*_*j*_(*t*).

Intercept-only (M-null), univariate, and multivariable (M-multi) models were evaluated independently for each ARG of clinical interest. Covariates were standardised (centred with their mean and scaled with their SD). To address multicollinearity, Pearson correlation coefficients were computed. For each exposome category (antibiotics, biocides, heavy metals, and anti-inflammatory drugs), covariates significantly correlated exclusively with a single sampling location were considered site-specific and excluded prior to aggregating overall exposome covariates (see Supplementary Text S2) ^52^. Pairwise correlations were then assessed among all covariates (Supplementary Figure S3). Each covariate was tested individually in a tested in the univariate analysis. Model performance was compared using Akaike Information Criterion (AIC) values between M-null and M-multi models. Models were fitted using restricted maximum likelihood (REML) as implemented in the lmer function of the lme4 R package (version 1.1.3.7). P-values for fixed effects were obtained using the lmerTest package. Genes *bla*_NDM_ and *mcr-1* had approximately two-thirds of observations below the detection limit and were therefore excluded from the mixed-effects model analysis due to insufficient quantifiable data for reliable parameter estimation (Supplementary Table S3).

In sensitivity analyses, we fitted an alternative model in which the response variable was defined as the change in relative ARG abundance between upstream and downstream sampling points, using the same covariates and random effect as in equation (1):

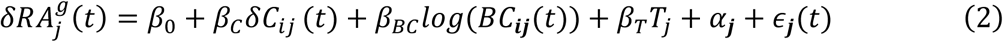

With *δRA_j_*^*g*^(*t*) represents the difference in relative abundance between upstream (*i)* and downstream (*j)* for a given gene *g* during week *t*. All other terms in the equation are defined in equation (1).

#### Spatio temporal dynamics in ESKAPEE

A mixed-effects model was used to analyse the relative abundance of ESKAPEE-associated genera (*Enterococcus*, *Staphylococcus*, *Klebsiella*, *Acinetobacter*, *Pseudomonas*, *Enterobacter*, and *Escherichia*) along the continuums over time and to identify associated factors while accounting for the dependence among sampling locations. Thus in this model, the response variable was the relative abundance of ESKAPEE-associated genera at each downstream sampling point and sampling date. The same covariate data was used as in equation (1) as well as a spatial random effect.

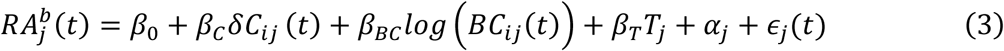

With *RA_j_*^*b*^(*t*) represents the relative abundance of a given bacteria *b* at downstream sampling point *j* during week *t*. All other variables in this equation are defined in equation (1).

All analyses were conducted in R (version 4.3.2).

## Acknowledgments

This work was supported by the French National Research Agency (ANR), grant number ANR-20-AMRB-0001-01.

