## Supplementary material for "From sewage to shoreline: Tracing antibiotic resistance gene trends through tropical island wastewater treatment pathways": -

### *Appendix 1*

### Supplementary methods

**Supplementary Table S1. Sampling point locations.**

| Sampling point | Name | Description | Lat, long |
|---|---|---|---|
| 1 | Hospital sewage | Hospital sewer before the water joins the municipal water system | 16.2374,<br>-61.5252 |
| 2 | Hospital: Influent | At the entrance of WWTP A (mixed hospital and municipal wastewater) | 16.2333,<br>-61.55 |

| | | | |
|---|---|---|---|
| 3 | Hospital: Effluent | At the exit of WWTP A | 16.2333, -61.55 |
| 4 | Hospital: Ocean | In the ocean after the exit of the WWTP A | 16.219913, -61.561047 |
| 5 | Domestic: Influent | At the entrance of WWTP B (municipal wastewater from the small town and poultry farms) | 16.284596, -61.631024 |
| 6 | Domestic: Effluent | At the exit of WWTP B | 16.284596, -61.631024 |
| 7 | Domestic: Mangrove | In the mangrove after the exit of WWTP B | 16.285261 -61.631099 |
| 8 | Hotel sewage | Wastewater coming from multiple hotels | 16.207970, -61.504663 |
| 9 | Touristic: Influent | At the entrance of WWTP C (mixed hotel and municipal wastewater) | 16.216156, -61.504035 |
| 10 | Touristic: Effluent | In the Mangrove after exit of WWTP C | 16.216156, -61.504035 |
| 11 | Touristic: Mangrove | In the Mangrove after the WWTP C | 16.214334 -61.508055 |

**Supplementary Text S1. Statistical Framework for Assessing Intra- and Inter-Filter Variability**

*Intra-Filter Variability*

To assess variability among the three technical replicates within each filter, we compared the distribution of CT values across all target genes. The underlying hypothesis was that replicates should exhibit consistent distributions. Statistical analysis was performed using the Wilcoxon signed-rank test, with p-values adjusted using the Benjamini-Hochberg method and a significance threshold of 0.05. If significant differences were detected, one or all replicates for the affected filter were removed. Specifically, if a single replicate exhibited a statistically significant deviation from the others, only that replicate was removed. Conversely, if all replicates were statistically different from one another, they were all excluded. After this filtering step, the relative abundance of genes was calculated, and the mean value for each filter was retained for further analysis.

*Inter-Filter Variability*

The next step was to assess the consistency of measurements across different filters. We analyzed differences in the distribution of mean relative abundances for all target genes between filters using a Kruskal-Wallis test. Dunn's post-hoc test was then applied to identify specific pairwise differences between filters, with p-values adjusted using the Benjamini-Hochberg method and a significance threshold of 0.05.

*Results Intra-Filter Variability*

No significant differences were observed between technical replicates within individual filters for both the hospital and Domestic continuums (Figure S1). However, in the touristic continuum, statistical differences were detected between technical replicates in the following cases: Filter 3 during week 7 of the touristic sample, Filter 2 during week 4 of the influent sample, and Filters 2 and 3 during week 4 of the effluent sample.

Based on these results, the following technical replicates were removed:

- Touristic continuum, week 7 – Touristic hotels sample, Filter 3, Replicate 2
- Touristic continuum, week 4 – Influent sample, Filter 2, Replicate 1
- Touristic continuum, week 4 – Effluent sample, Filter 2, all replicates
- Touristic continuum, week 4 – Effluent sample, Filter 3, Replicate

*Results Inter-Filter Variability*

In the hospital continuum, during the third campaign (week 9), Filter 3 exhibited a significantly higher relative abundance compared to the other two filters and was therefore removed from the analysis (Figure S2). *(Note: Data from Campaign 1 had already been excluded.)*

In the Domestic continuum, statistical differences were observed between samples from weeks 4 and 10 of the influent sample (Figure S2).

In the touristic continuum, Filters 2 and 3 in week 4 (influent) displayed significantly lower relative abundance values than Filter 1. (Figure S2). While examining only week 4, Filters 2 and 3 appeared more similar to each other, suggesting that Filter 1 might be an outlier. However, when considering data from all filters across multiple time points, the overall trend indicated that Filters 2 and 3 were the likely outliers. Statistical differences were also observed between samples from week 4 of the effluent sample point.

Based on these results, the following technical replicates were removed:

- Hospital continuum, week 9 – Effluent sample, Filter 3, all replicates (Inter-filter variability)
- Touristic continuum, week 7 – Hotel sewage sample, Filter 3, Replicate 2 (Intra-filter variability)
- Touristic continuum, week 4 – Influent sample, Filters 2 and 3, all replicates (Intra-filter + Inter-filter variability)

**Supplementary Figure S1.Cycle threshold distribution of all ARGSs and MGEs before (influent) and after (effluent) wastewater treatment for each technical replicate.** The horizontal numbers represent the sampling weeks. Red dots indicate the cycle threshold of the 16S gene. Asterisks indicate significant differences between technical replicates (* $p<0.05$, ** $p<0.01$, *** $p<0.001$, **** $p<0.0001$).  *Here "Non-touristic continuum" refers to the domestic continuum.

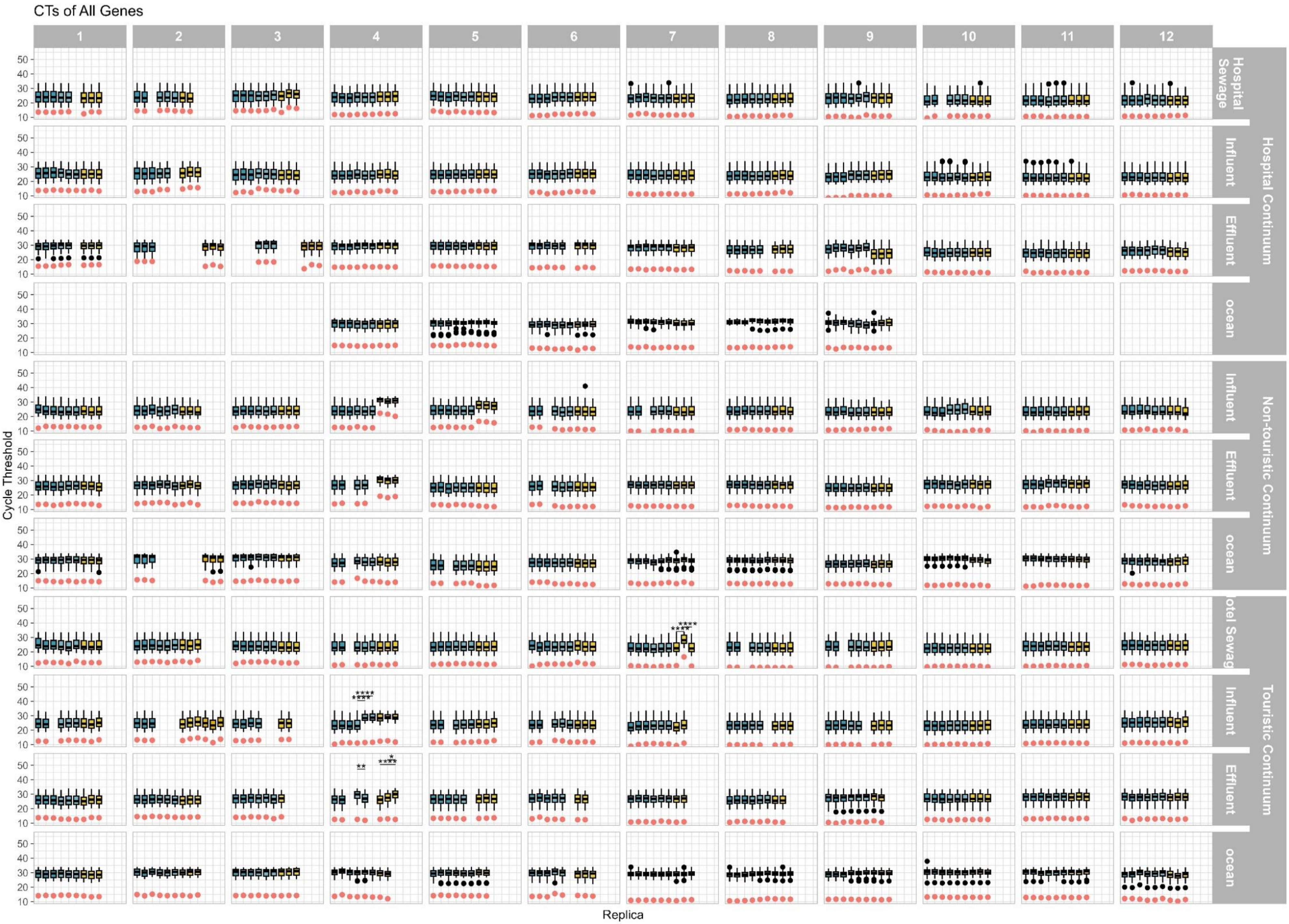
CTs of All Genes
1
2
3
4
5
6
7
8
9
10
11
12
Hospital Sewage
Influent
Effluent
ocean
Hospital Continuum
Influent
Effluent
ocean
Non-touristic Continuum
otel Sewag
Influent
Effluent
ocean
Touristic Continuum
Cycle Threshold
Replica
50
40
30
20
10

**Supplementary Figure S2. Relative abundance of all ARGSs and MGEs before (influent) and after (effluent) wastewater treatment for all filters.** The horizontal numbers represent the sampling weeks. Asterisks indicate significant differences between filters (* p<0.05, ** p<0.01, *** p<0.001, **** p<0.0001). *Here "Non-touristic continuum" refers to the domestic continuum.

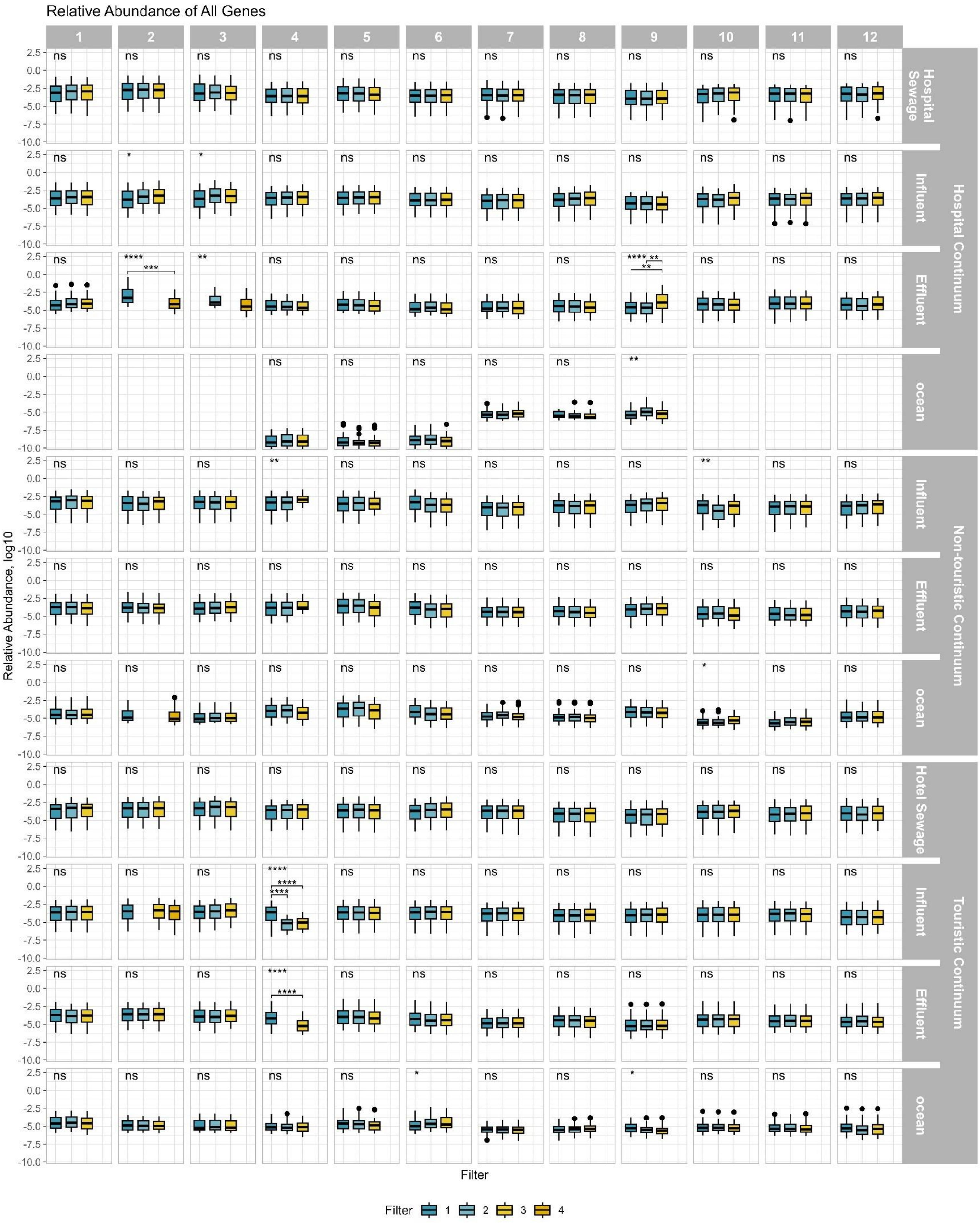
Relative Abundance of All Genes
1
2
3
4
5
6
7
8
9
10
11
12
Hospital Sewage
Influent
Effluent
ocean
Hospital Continuum
Influent
Effluent
ocean
Non-touristic Continuum
Hotel Sewage
Influent
Effluent
ocean
Touristic Continuum
Relative Abundance, log10
Filter
Filter 1 2 3 4

**Supplementary Table S2. All collected data.**

Resistome and exposome data were collected at all sampling points. Climate data is only collected once per continuum at the WWTP location. Resistome: 80 targeted antibiotic-resistant genes (ARGs) grouped into 16 resistance classes and 12 mobile genetic elements (MGEs), including three integrons. Exposome: antibiotics, biocides, non-steroidal anti-inflammatory drugs, and heavy metals. 16 clinically relevant ARGs and MGEs are in bold.

| RESISTOME | Genes conferring resistance to: | ARGs and MGEs |
|---|---|---|
| | Chloramphenicol | catB3, cat, cmlA1 |
| | Aminoglycosides | aac(3)-lid, aadE, aac(6')-lb, aadA, aac(6')-llA, spc, aph(3')-III, aph(2)-l(de), aph(2)-lb, aadB, AAC(3)-lb, strB, aac(6)-aph(2) |
| | Bacitracin | bacA_2, bacA_1 |
| | Beta-lactams | cepA, cblA, cfxA, blaTEM, blaIMP, blaAmpC, blaDHA, blaCMY-2, blaACC, blaSHV, blaVIM, blaKPC, blaIMI, blaBIC-1, blaGES, blaNDM, blaOXA, blaCTX-M, blaPER-1 |
| | Macrolides | ermF, ermB, ermC, ermY, mfsA, mefA_10, macB, ermX |
| | Efflux pumps | acrA, mdtO, mdtL, mdtF, tolC |
| | Quinolones | qnrA, qnrB, qnrS, qnrC |
| | Heavy metals | cusF, copA, copD, cadA, merA, czcA |
| | QACs | qacA, qacC, qacE |
| | Vancomycin | vanA, vanB |
| | Tetracyclines | tetQ, tetW, tetM, tetO, tetB |
| | Polymyxin | arnA, mcr-1 |
| | Sulfonamides | sul1, sulA |
| | Methicillin | mecA |
| | Trimethoprim | dfrA27, dfrF, dfrB1 |
| | Streptogramins | vat(A), vatB |

| | | |
|---|---|---|
| | Mobile genetic elements (MGEs): | ISSW1, ISS1N, IS6_IS6100, Tp614, IS613, IS6 group, tn3_tnpA, ISEc9, inc-P1, integrons (intI1, intI2, intI3) |
| EXPOSOME | Compound | Name |
| | Antibiotic (ng/L) | Azithromycin, Ciprofloxacin, Erythromycin, Ofloxacine, Pyrimethamine, Sulfaguanidine, Sulfamethoxazole, Sulfathiazole, Trimethoprime |
| | Biocide (ng/L) | Acephate, Albendazole, Flubendazole, Levamisole, Pyrantel pamoate, Sulfoxaflor, Tebuconazole, Thiabendazole |
| | Anti-inflammatory (ng/L) | Diclofenac, Ibuprofen, Ketoprofen |
| | Heavy metal (µg/L) | Cr, Zn, Cu, As, Cd, Gd, Hg, Pb |
| CLIMATE DATA | Type of data | Measurement |
| | Climate parameters | Max temperature (°C),<br>Min temperature (°C),<br>Water temperature (°C),<br>Precipitation (mm) D-2,<br>Precipitation (mm) D-1,<br>Precipitation (mm) D0<br>(D-2 = two days before the sampling, D-1= day before the sampling, D0 = day of the sampling) |

**Supplementary Text S2. Selection and Construction of Exposome Covariates.**

*Step 1 – Identification of site-specific covariates*

For each exposome category (antibiotics, biocides, heavy metals, and anti-inflammatory drugs), we assessed the association between individual compounds and sampling-point indicator variables. Compounds significantly associated with only one sampling point were considered to exhibit a site-specific signature and were therefore excluded from the computation of overall category concentrations, as they primarily reflected local contamination sources rather than broader exposome patterns.

For antibiotics, Azithromycin and Erythromycin (macrolides) and Sulfaguanidine (sulfonamides) were retained. In contrast, Ciprofloxacin, Ofloxacin, Sulfamethoxazole, and Trimethoprim were exclusively associated with the Hospital Sewage sampling point ($p < 0.005$) and were therefore classified as site-specific. Sulfathiazole was similarly identified as site-specific due to its exclusive association with the Domestic Mangrove site.

Among biocides, Albendazole was significantly associated with Hospital Sewage, Sulfoxaflor with Hospital Influent, and Pyrimethamine with Hospital Ocean; these compounds were therefore considered site-specific. The remaining biocides retained for further analyses were Flubendazole, Levamisole, Pyrantel pamoate, Tebuconazole, and Thiabendazole.

For heavy metals, Copper (Cu), Gadolinium (Gd), and Mercury (Hg) displayed site-specific associations with Hospital Ocean, Hospital Sewage, and Domestic Influent, respectively, and were excluded. The heavy metals retained for subsequent analyses were Arsenic (As), Cadmium (Cd), Chromium (Cr), Lead (Pb), and Zinc (Zn).

No anti-inflammatory drugs showed exclusive associations with a single sampling point; consequently, all anti-inflammatory compounds were retained.

*Step 2 – Computation of category-specific overall concentrations*

After removal of site-specific compounds, category-specific overall concentrations were calculated using the remaining covariates within each exposome category. This approach ensured that the resulting metrics captured generalized exposure patterns across sampling locations while minimizing the influence of compounds reflecting localized contamination sources.

**Supplementary Figure S3. Correlogram for covariables data used in the mixed effect model.**

Shown here are the Pearson correlation coefficient for each pair of variables (r). Blue colors represent positive correlation between variables. Red colors represent negative correlation. Only significant correlation coefficients (p-value < 5%) are shown in the correlogram.

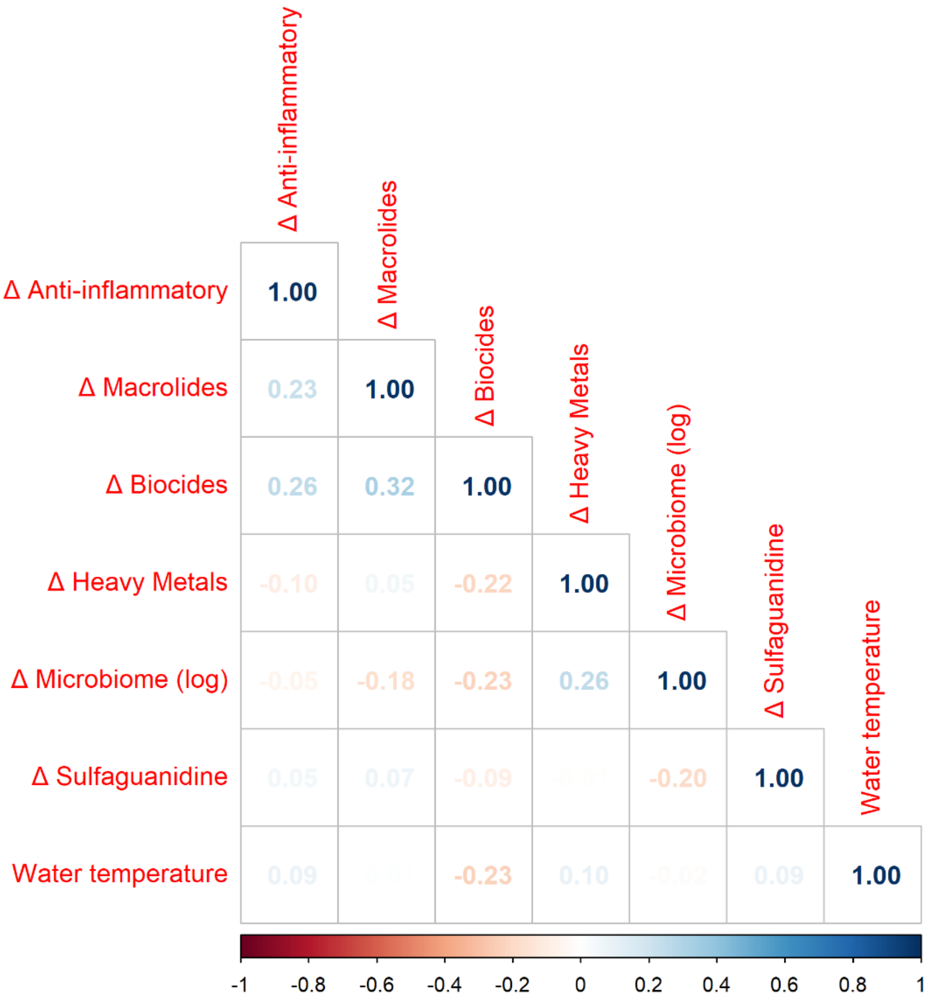

**Supplementary Table S3. Number of observations included in the mixed-effect model for each clinically relevant gene.**

All sampled points ($n = 84$) were included, except observations excluded because the relative abundance was below the limit of detection or because one or more covariates were missing. For example, microbiome dissimilarity values for the last week in the Domestic mangrove were unavailable.

| Gene | Number of observations |
|---|---|
| *aac(6')-I B* | 82 |

| | |
|---|---|
| *aph(3')-III* | 77 |
| *blaCTX-M* | 67 |
| *blaKPC* | 66 |
| *blaNDM* | 28 |
| *blaOXA* | 72 |
| *blaSHV* | 69 |
| *blaTEM* | 83 |
| *blaVIM* | 60 |
| *ermB* | 83 |
| *intI1* | 75 |
| *mcr-1* | 29 |
| *mecA* | 58 |
| *qnrS* | 83 |
| *sul1* | 75 |
| *tetM* | 83 |

### Supplementary data description

**Supplementary Figure S4. Total relative abundance per sampling point.**

Each dot represents a sampling week. *Here “Non-touristic” refers to the domestic continuum.

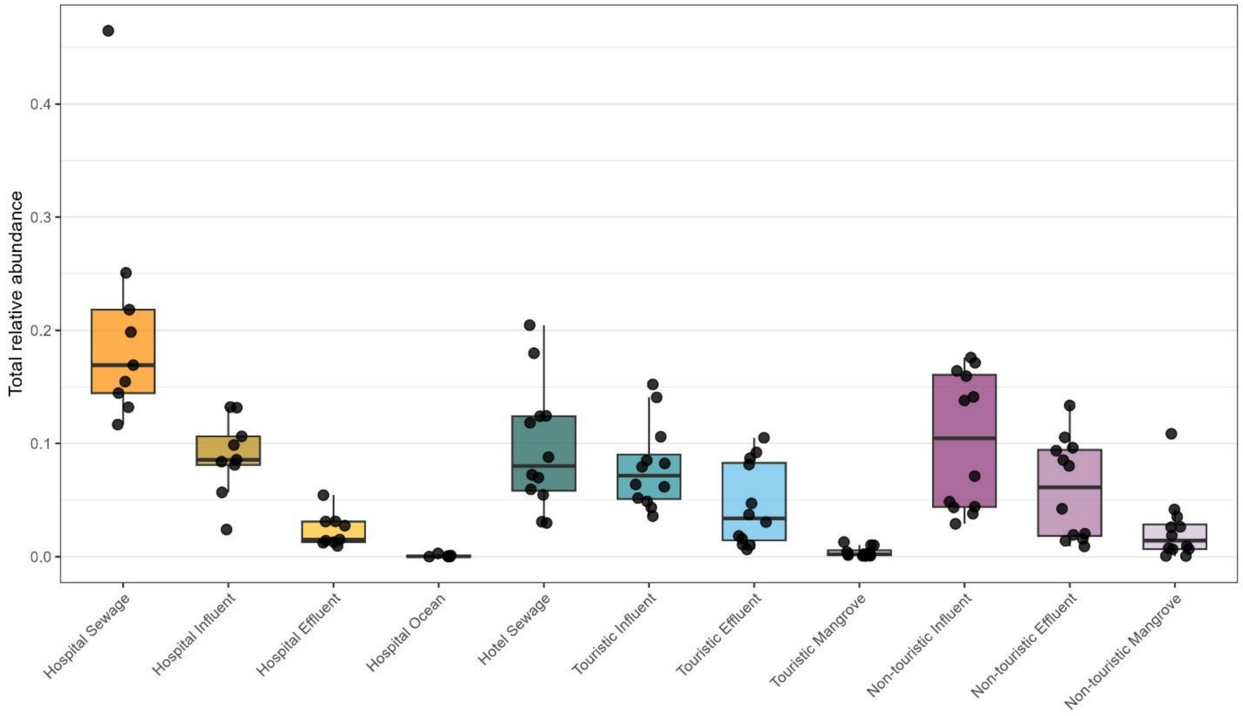

**Supplementary Figure S5. Antibiotic boxplot per sampling point.**

Concentrations are given in ng/L. *Here “Non-touristic” refers to the domestic continuum.

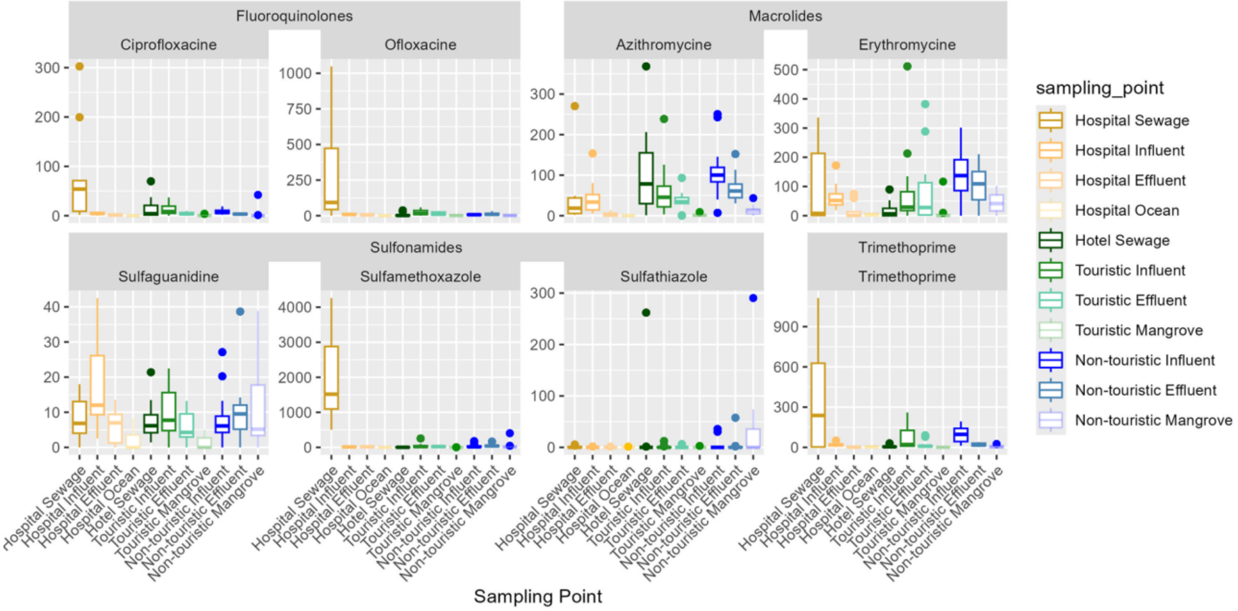

**Supplementary Figure S6. Biocide boxplot per sampling point.**

Concentrations are given in ng/L. *Here “Non-touristic” refers to the domestic continuum.

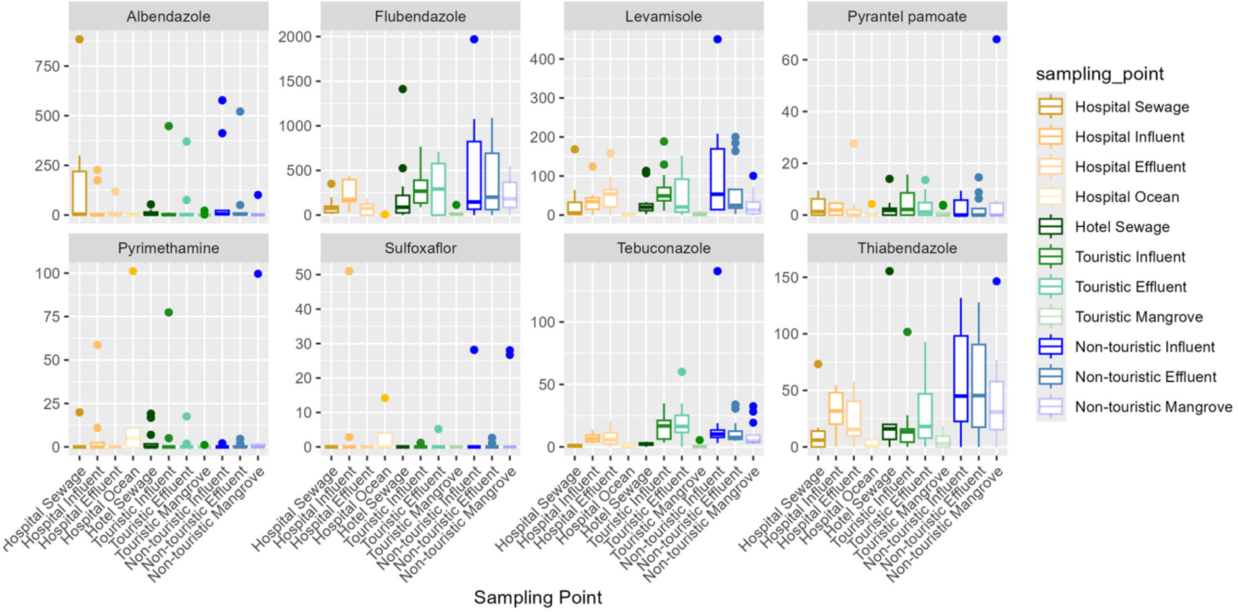

**Supplementary Figure S7. Heavy metals boxplot per sampling point.**

Concentrations are given in µg/L. *Here “Non-touristic” refers to the domestic continuum.

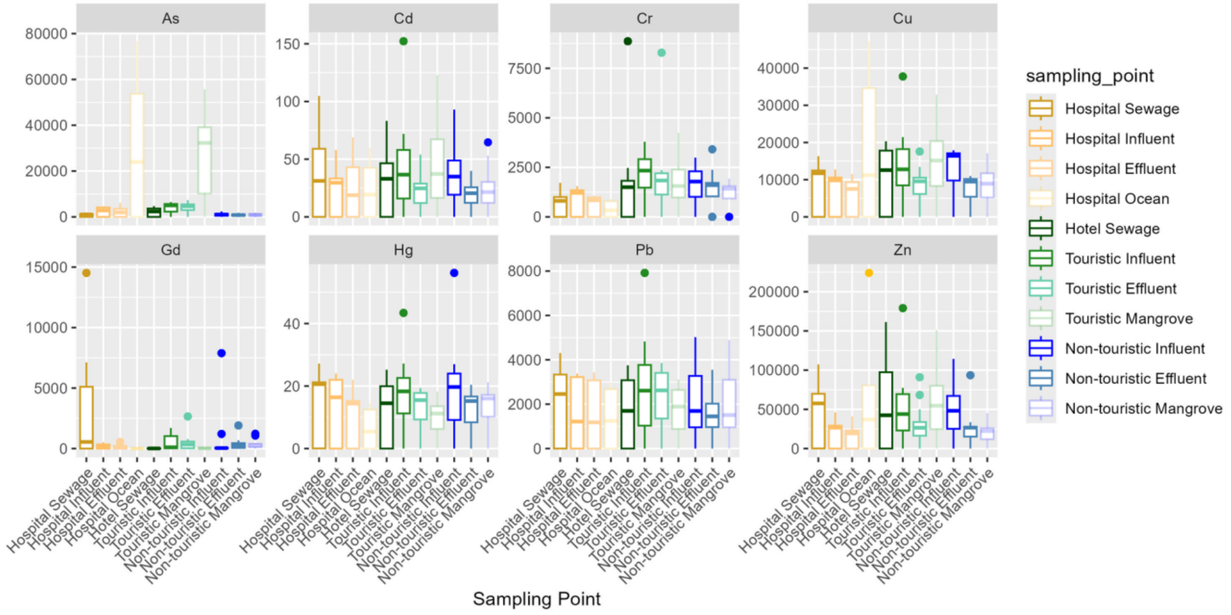

**Supplementary Figure S8. Non-steroidal anti-inflammatory drugs concentration per sampling point.**

Concentrations are given in ng/L. *Here "Non-touristic" refers to the domestic continuum.

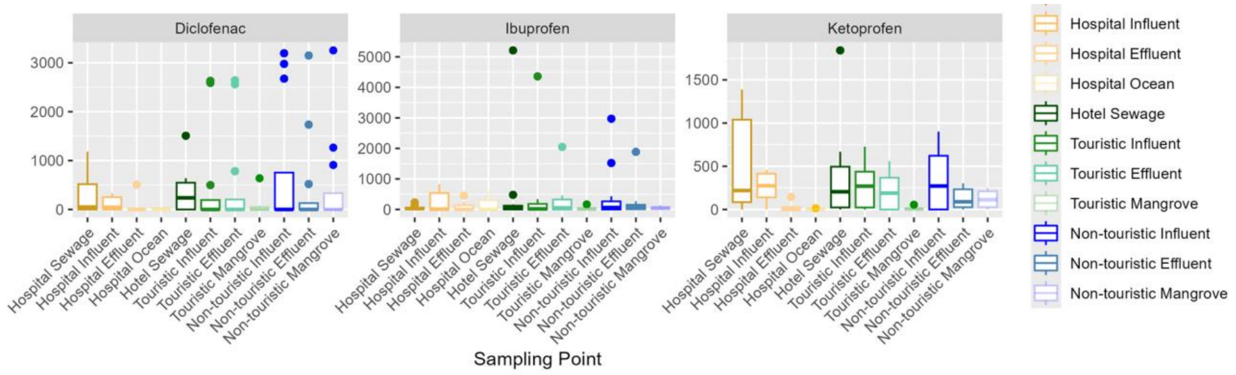

### Supplementary results

**Supplementary Table S4. Genera identified as members of the core community for each sampling point.**

The first column shows the genera. The second to last column show the average relative abundance for each sampling point; core genera are indicated by bold squares. Color of the square is only for visualization purposes; as higher average relative abundance is indicated by a more red color. *Here “Non-touristic” refers to the domestic continuum.

| | Hospital Sewage | Hotel Sewage | Hospital Influent | Touristic Influent | Non-touristic Influent | Hospital Effluent | Touristic Effluent | Non-touristic Effluent | Hospital Ocean | Touristic Mangrove | Non-touristic Mangrove |
|---|---|---|---|---|---|---|---|---|---|---|---|
| Arcobacter | 7.68% | 9.64% | 20.21% | 10.58% | 10.83% | 9.65% | 7.71% | 12.67% | 1.01% | 11.67% | 5.66% |
| Prochlorococcus | 0.00% | 0.03% | 0.00% | 10.28% | 0.00% | 0.61% | 0.02% | 0.12% | 59.94% | 28.01% | 0.01% |
| Streptococcus | 7.10% | 12.30% | 5.16% | 12.41% | 15.86% | 2.07% | 1.90% | 2.45% | 0.49% | 2.19% | 4.82% |
| Aeromonas | 10.54% | 3.03% | 7.08% | 4.34% | 5.74% | 5.14% | 1.65% | 5.46% | 0.42% | 0.24% | 3.44% |
| Bacteroides | 3.86% | 6.02% | 6.25% | 3.16% | 4.87% | 6.46% | 1.62% | 6.84% | 0.39% | 0.37% | 3.38% |
| Pseudomonas | 2.85% | 4.62% | 5.33% | 2.81% | 4.82% | 3.71% | 4.00% | 6.75% | 0.30% | 0.88% | 4.06% |
| Acinetobacter | 1.56% | 4.87% | 5.13% | 7.43% | 8.71% | 4.31% | 1.77% | 1.56% | 0.09% | 0.61% | 2.16% |
| Flavobacterium | 0.50% | 0.06% | 0.28% | 1.46% | 0.15% | 4.97% | 7.35% | 15.96% | 0.17% | 1.09% | 4.09% |
| Cloacibacterium | 1.55% | 1.93% | 5.35% | 6.94% | 6.93% | 2.42% | 3.04% | 3.48% | 0.17% | 0.44% | 3.18% |
| Parabacteroides | 0.86% | 1.73% | 4.33% | 4.11% | 3.36% | 2.18% | 1.68% | 2.74% | 0.09% | 0.59% | 1.96% |
| Brachymonas | 0.01% | 0.07% | 0.10% | 0.48% | 0.07% | 0.03% | 18.73% | 0.05% | 1.50% | 2.38% | 0.05% |
| Thauera | 0.10% | 0.19% | 0.33% | 0.38% | 0.16% | 0.17% | 15.51% | 0.24% | 0.50% | 2.06% | 0.17% |
| Diaphorobacter | 7.86% | 0.90% | 1.69% | 1.92% | 2.75% | 1.25% | 0.57% | 0.97% | 0.10% | 0.44% | 0.62% |
| Prevotella | 3.04% | 3.62% | 1.06% | 0.70% | 1.87% | 0.80% | 0.16% | 1.74% | 0.19% | 0.06% | 3.47% |
| Blautia | 2.63% | 5.70% | 1.32% | 1.52% | 2.77% | 0.60% | 0.25% | 0.58% | 0.06% | 0.14% | 1.05% |
| Aquaspirillum | 0.75% | 1.42% | 2.65% | 2.04% | 0.82% | 2.41% | 1.57% | 2.98% | 0.08% | 1.22% | 0.56% |
| Thiothrix | 0.12% | 0.11% | 3.41% | 1.08% | 0.42% | 1.47% | 1.37% | 0.29% | 0.09% | 6.93% | 0.56% |
| Clostridium | 1.95% | 1.98% | 0.83% | 0.79% | 0.93% | 2.39% | 0.99% | 1.45% | 0.06% | 0.26% | 4.11% |
| Faecalibacterium | 5.15% | 4.61% | 0.84% | 1.24% | 1.17% | 0.43% | 0.29% | 0.09% | 0.06% | 0.25% | 0.80% |
| Eubacterium | 3.69% | 3.12% | 0.92% | 0.90% | 2.34% | 0.49% | 0.24% | 0.38% | 0.04% | 0.22% | 0.58% |
| Sulfurimonas | 0.01% | 0.00% | 0.40% | 0.59% | 0.00% | 0.14% | 3.15% | 0.00% | 0.18% | 5.13% | 0.12% |
| Sulfuricurvum | 0.01% | 0.07% | 0.31% | 0.12% | 0.04% | 0.27% | 1.58% | 0.06% | 0.00% | 0.10% | 4.45% |

**Supplementary Table S5. Alpha diversity Chaos 1 richness comparison between sampling points.**

Wilcoxon test with Bonferroni adjustment no paired samples

| Continuum | Sampling point 1 | Sampling point 2 | p.adj | Significance |
|---|---|---|---|---|
| Hospital | Hospital Sewage | Hospital Influent | 1.000000 | ns |
| | Hospital Influent | Hospital Effluent | 0.157000 | ns |
| | Hospital Effluent | Hospital Ocean | 0.026000 | * |

| | | | | |
|---|---|---|---|---|
| Domestic | Domestic Influent | Domestic Effluent | 1.000000 | ns |
| | Domestic Effluent | Domestic Mangrove | 1.000000 | ns |
| Touristic | Hotel Sewage | Touristic Influent | 1.000000 | ns |
| | Touristic Influent | Touristic Effluent | 1.000000 | ns |
| | Touristic Effluent | Touristic Mangrove | 0.012000 | * |

**Supplementary Table S6. Alpha diversity Shannon diversity comparison between sampling points.**

Shannon diversity. Wilcoxon test with Bonferroni adjustment no paired samples

| Continuum | Sampling point 1 | Sampling point 2 | p.adj | Significance |
|---|---|---|---|---|
| Hospital | Hospital Sewage | Hospital Influent | 1.000000 | ns |
| | Hospital Influent | Hospital Effluent | 1.000000 | ns |
| | Hospital Effluent | Hospital Ocean | 0.000648 | *** |
| Domestic | Domestic Influent | Domestic Effluent | 1.000000 | ns |
| | Domestic Effluent | Domestic Mangrove | 0.272000 | ns |
| Touristic | Hotel Sewage | Touristic Influent | 1.000000 | ns |
| | Touristic Influent | Touristic Effluent | 0.087000 | ns |
| | Touristic Effluent | Touristic Mangrove | 1.000000 | ns |

**Supplementary Figure S9. Top 10 genera contributing to the first two principal coordinates of the PcoA using the microbiome Bray–Curtis dissimilarity matrix.**

A. Variables contributing to the first principal coordinate. B. Variables contributing to the second principal coordinate.

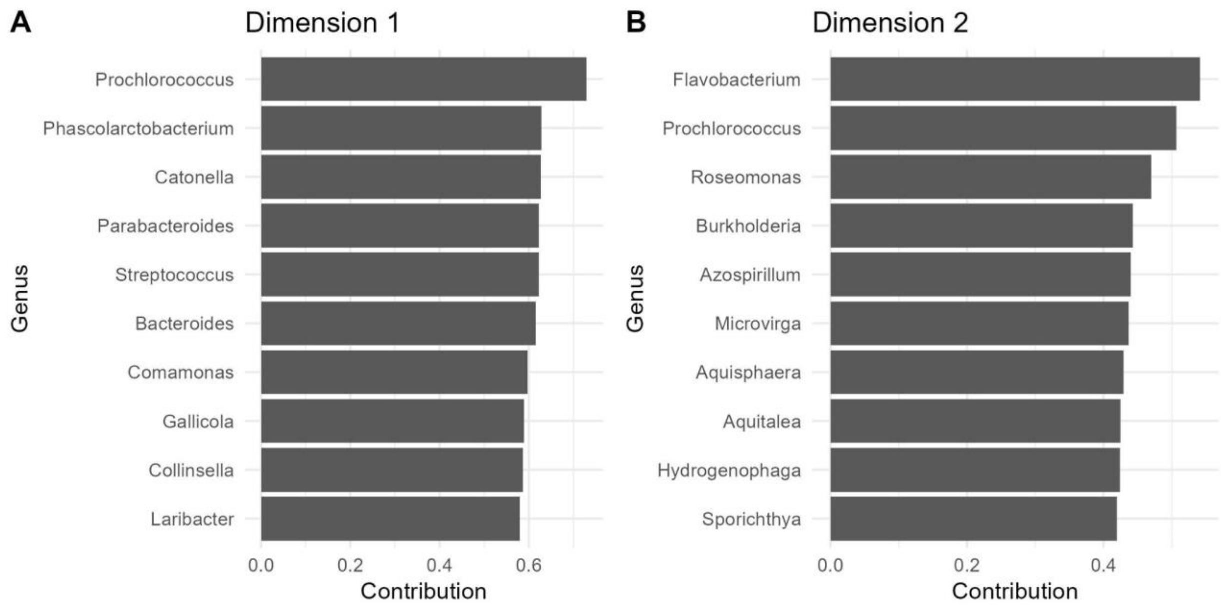

**Supplementary Figure S10. Top 10 ARGs contributing to the first two principal coordinates of the PcoA using the resistome Bray–Curtis dissimilarity matrix.** A. Variables contributing to the first principal coordinate. B. Variables contributing to the second principal coordinate.

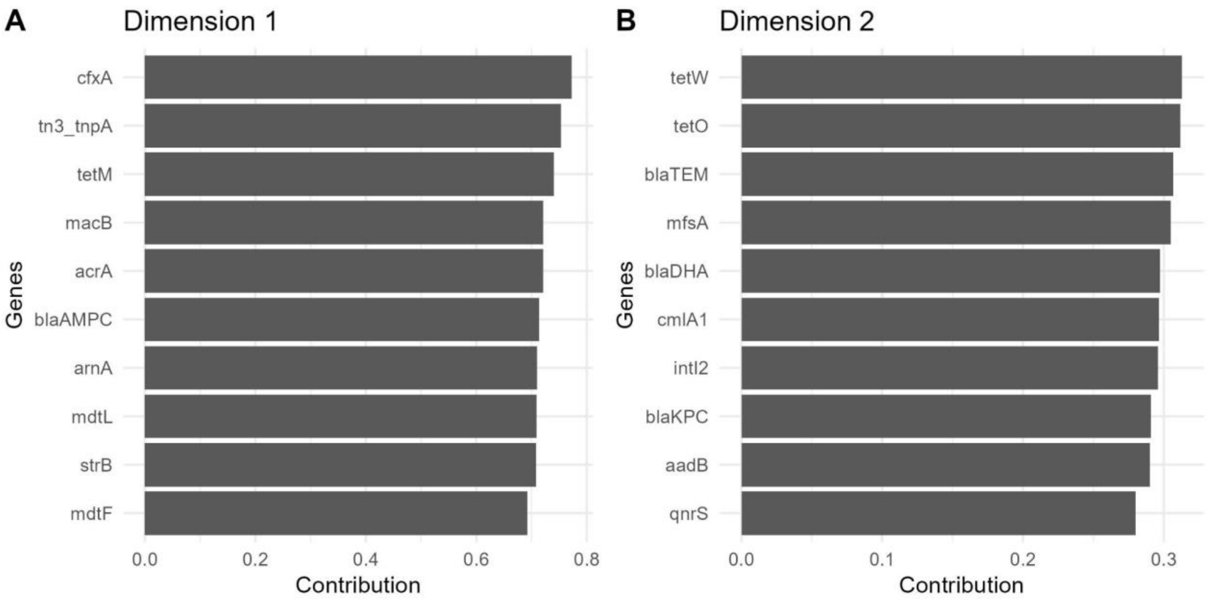

**Supplementary Figure S11. Resistome clustering of all ARGs and MGEs.**
Clustered heatmap of all genes. Colors in the heatmap represent log_10 transformed relative abundance. Columns represent each sample collected and are clustered based on the full resistome Bray-Curtis dissimilarity matrix. Rows represent each ARG and are clustered based on Pearson correlation using complete-linkage. Annotation colors for columns (samples) are based on water temperature, water type (sewage, influent, effluent and ocean/mangrove) and continuum. Annotation colors for row (genes) are based on the resistance type.

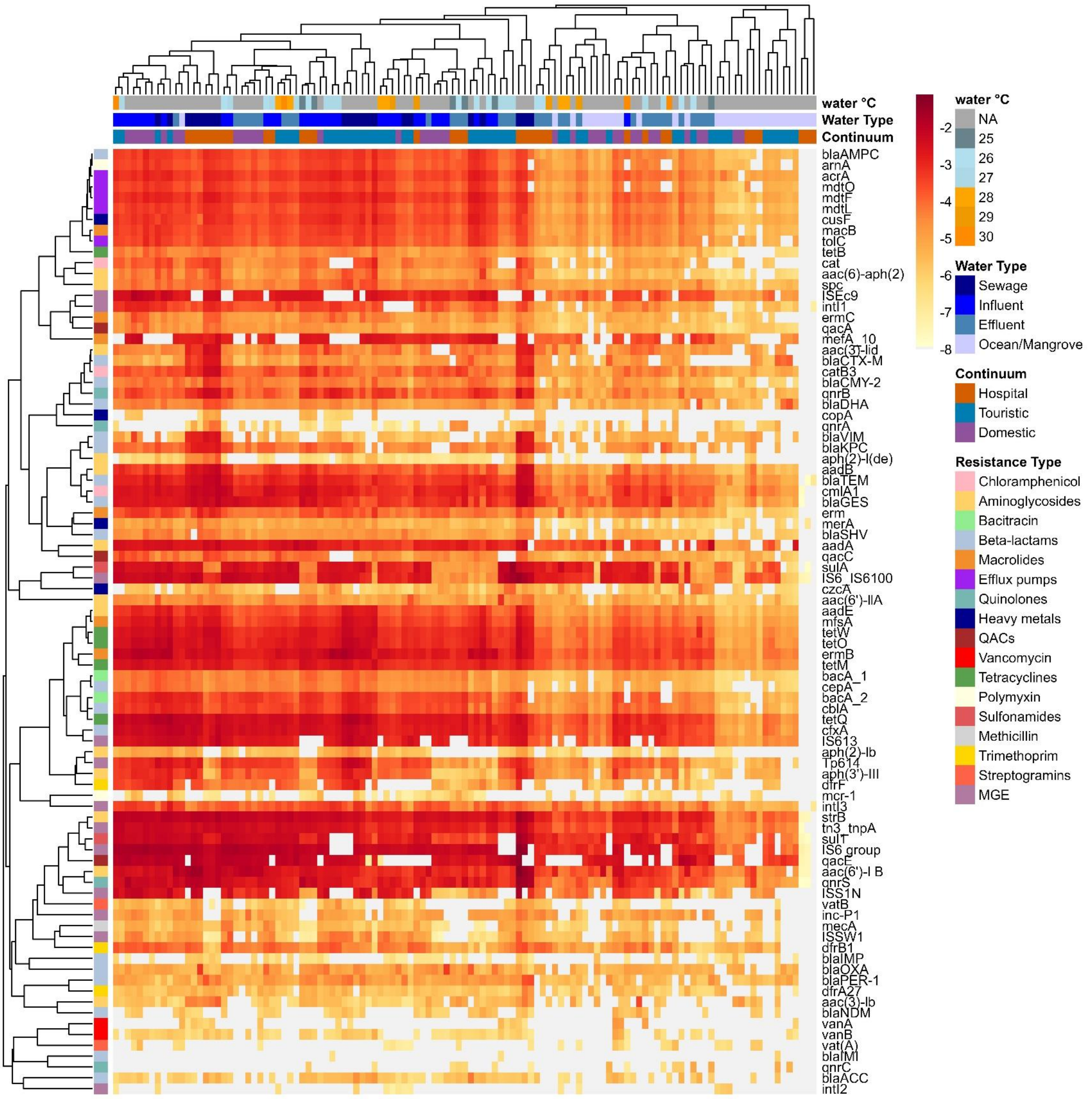

water °C
Water Type
Continuum
blaAMPC
arnA
acrA
mdtO
mdtF
mdtL
cusF
macB
tolC
tetB
cat
aac(6)-aph(2)
spc
ISEc9
intI1
ermC
qacA
mefA_10
aac(3)-iid
blaCTX-M
catB3
blaCMY-2
qnrB
blaDHA
copA
qnrA
blaVIM
blaKPC
aph(2')-I(de)
aadB
blaTEM
cmlA1
blaGES
erm
merA
blaSHV
aadA
qacC
sulA
IS6_IS6100
czcA
aac(6')-IIA
aadE
mfsA
tetW
tetO
ermB
tetM
bacA_1
cepA
bacA_2
cblA
tetQ
cfxA
IS613
aph(2)-Ib
Tp614
aph(3')-III
dfrF
mcr-1
intI3
strB
tn3_tnpA
sulT
IS6 group
qacE
aac(6')-I B
qnrS
ISS1N
vatB
inc-P1
mecA
ISSW1
dfrB1
blaIMP
blaOXA
blaPER-1
dfrA27
aac(3)-Ib
blaNDM
vanA
vanB
vat(A)
blaIMI
qnrC
blaACC
intI2
-2
-3
-4
-5
-6
-7
-8
water °C
NA
25
26
27
28
29
30
Water Type
Sewage
Influent
Effluent
Ocean/Mangrove
Continuum
Hospital
Touristic
Domestic
Resistance Type
Chloramphenicol
Aminoglycosides
Bacitracin
Beta-lactams
Macrolides
Efflux pumps
Quinolones
Heavy metals
QACs
Vancomycin
Tetracyclines
Polymyxin
Sulfonamides
Methicillin
Trimethoprim
Streptogramins
MGE

### Supplementary model's results

**Supplementary Figure S12. Observed vs fitted values by mixed-effect model, for each gene of clinical interest.** Red points are the observed relative abundance by sampling point- week of sampling with corresponding 95% confidence intervals around the prediction**.**

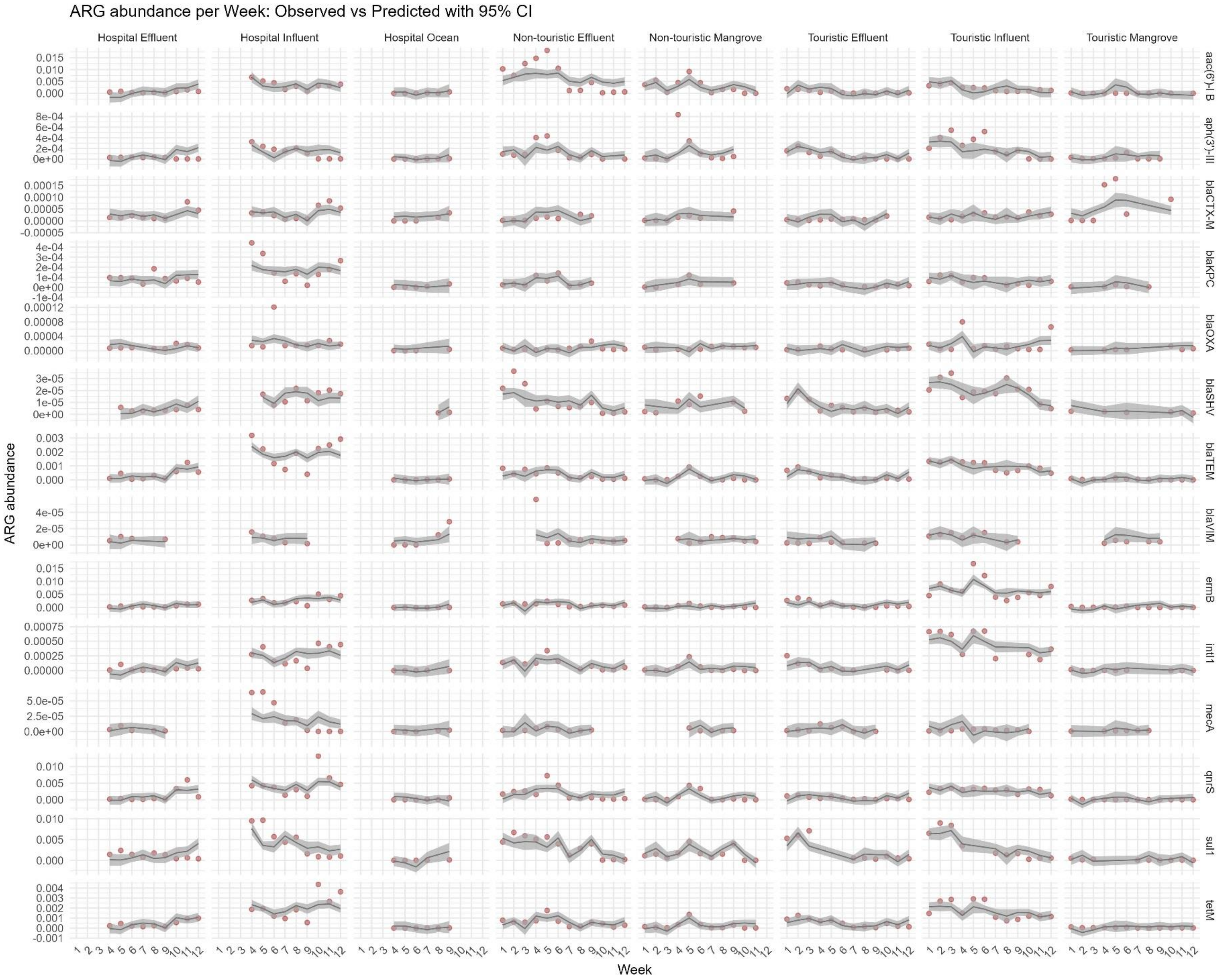

**Supplementary Table S7. Univariate results between heavy metals and each gene of clinical interest.**

Estimated coefficients from univariate analyses, using the mixed-effect model. Coefficients associated with a p-value < 5% are highlighted in bold. Cr: Chromium, Zn: Zinc; As: Arsenic; Cd: Cadmium; Pb: Lead.

| Gene | ΔCr | ΔZn | ΔAs | ΔCd | ΔPb |
|---|---|---|---|---|---|
| *aac(6')-I B* | -5.8e-05<br>[-6.8e-04, 5.6e-04] | 4.4e-04<br>[-2.3e-04, 1.1e-03] | -2.3e-04<br>[-1.0e-03, 5.6e-04] | 3.1e-04<br>[-3.6e-04, 9.8e-04] | 1.7e-05<br>[-6.5e-04, 6.8e-04] |
| *aph(3')-III* | 1.7e-05<br>[-1.6e-05, 5.1e-05] | 8.4e-06<br>[-2.8e-05, 4.4e-05] | -1.9e-05<br>[-5.8e-05, 1.9e-05] | 1.1e-05<br>[-2.4e-05, 4.6e-05] | 2.3e-05<br>[-1.1e-05, 5.8e-05] |
| *blaCTX-M* | -3.0e-06<br>[-1.0e-05, 4.3e-06] | 5.3e-06<br>[-2.6e-06, 1.3e-05] | **2.0e-05**<br>**[1.3e-05, 2.8e-05]** | 5.9e-06<br>[-2.0e-06, 1.4e-05] | -3.4e-06<br>[-1.1e-05, 4.5e-06] |
| *blaKPC* | -1.8e-06<br>[-1.6e-05, 1.3e-05] | 6.5e-06<br>[-9.9e-06, 2.3e-05] | -3.7e-06<br>[-2.3e-05, 1.5e-05] | -5.9e-07<br>[-1.6e-05, 1.5e-05] | 1.9e-06<br>[-1.4e-05, 1.8e-05] |
| *blaOXA* | 8.9e-07<br>[-3.1e-06, 4.9e-06] | 1.6e-06<br>[-2.6e-06, 5.8e-06] | -5.6e-07<br>[-5.3e-06, 4.2e-06] | 2.1e-06<br>[-2.2e-06, 6.3e-06] | 2.7e-06<br>[-1.5e-06, 6.9e-06] |
| *blaSHV* | 9.4e-08<br>[-1.6e-06, 1.8e-06] | 8.9e-07<br>[-9.3e-07, 2.7e-06] | -9.5e-07<br>[-3.3e-06, 1.3e-06] | 1.6e-06<br>[-2.1e-07, 3.4e-06] | 3.0e-07<br>[-1.5e-06, 2.1e-06] |
| *blaTEM* | -5.5e-06<br>[-9.6e-05, 8.5e-05] | 5.9e-05<br>[-4.1e-05, 1.6e-04] | -1.9e-05<br>[-1.4e-04, 1.0e-04] | 2.0e-05<br>[-8.0e-05, 1.2e-04] | 1.1e-05<br>[-8.8e-05, 1.1e-04] |
| *blaVIM* | -2.1e-07<br>[-2.3e-06, 1.8e-06] | 1.5e-06<br>[-5.3e-07, 3.5e-06] | 7.2e-07<br>[-1.3e-06, 2.8e-06] | 3.3e-07<br>[-1.7e-06, 2.4e-06] | -2.9e-07<br>[-2.3e-06, 1.8e-06] |
| *ermB* | 3.3e-04<br>[-4.4e-05, 7.0e-04] | 3.3e-04<br>[-8.9e-05, 7.4e-04] | -2.2e-05<br>[-5.3e-04, 4.9e-04] | 1.1e-04<br>[-3.1e-04, 5.3e-04] | **7.1e-04**<br>**[3.3e-04, 1.1e-03]** |
| *intI1* | 1.0e-05<br>[-1.5e-05, 3.6e-05] | 2.1e-05<br>[-6.9e-06, 5.0e-05] | -3.9e-06<br>[-3.6e-05, 2.8e-05] | 5.8e-06<br>[-2.3e-05, 3.5e-05] | 7.6e-06<br>[-2.0e-05, 3.5e-05] |
| *mecA* | 1.2e-07<br>[-2.9e-06, 3.2e-06] | 1.8e-06<br>[-1.5e-06, 5.0e-06] | -9.1e-07<br>[-4.6e-06, 2.8e-06] | 1.5e-06<br>[-1.8e-06, 4.8e-06] | 1.5e-06<br>[-1.8e-06, 4.7e-06] |
| *qnrS* | -7.6e-05<br>[-4.5e-04, 3.0e-04] | 9.0e-05<br>[-3.3e-04, 5.0e-04] | -1.1e-04<br>[-6.0e-04, 3.8e-04] | -1.8e-04<br>[-5.9e-04, 2.3e-04] | -2.4e-04<br>[-6.4e-04, 1.7e-04] |
| *sul1* | 2.6e-04<br>[-2.8e-04, 8.1e-04] | 4.9e-04<br>[-7.9e-05, 1.1e-03] | -3.9e-04<br>[-1.0e-03, 2.3e-04] | 3.2e-04<br>[-2.6e-04, 8.9e-04] | 4.3e-04<br>[-1.4e-04, 1.0e-03] |

| Gene | ΔCr | ΔZn | ΔAs | ΔCd | ΔPb |
|---|---|---|---|---|---|
| *tetM* | 3.6e-05 [-9.6e-05, 1.7e-04] | 4.6e-05 [-1.0e-04, 1.9e-04] | -3.0e-05 [-2.1e-04, 1.5e-04] | -6.1e-05 [-2.1e-04, 8.5e-05] | -1.5e-05 [-1.6e-04, 1.3e-04] |

**Supplementary Table S8. Univariate results between climate variables and each gene of clinical interest.**

Estimated coefficients from univariate analyses, using the mixed-effect model.
Coefficients associated with a p-value < 5% are highlighted in bold.

| Gene | min °C | max °C | water | Precipitation D2 | Precipitation D1 | Precipitation D0 |
|---|---|---|---|---|---|---|
| aac(6')-l B | -4.7e-05 [-6.6e-04, 5.7e-04] | -2.3e-04 [-8.4e-04, 3.8e-04] | 2.6e-05 [-5.9e-04, 6.5e-04] | -7.1e-06 [-6.3e-04, 6.1e-04] | -2.2e-04 [-8.5e-04, 4.2e-04] | -3.8e-04 [-1.1e-03, 2.9e-04] |
| aph(3')-III | -2.4e-05 [-5.7e-05, 8.6e-06] | -7.9e-06 [-4.1e-05, 2.6e-05] | -1.0e-05 [-4.4e-05, 2.4e-05] | -2.6e-06 [-3.6e-05, 3.1e-05] | -3.7e-06 [-3.8e-05, 3.0e-05] | -2.9e-05 [-6.4e-05, 5.5e-06] |
| blaCTX-M | **-1.1e-05 [-1.8e-05, -3.9e-06]** | **-1.2e-05 [-1.9e-05, -5.7e-06]** | **-1.1e-05 [-1.8e-05, -4.5e-06]** | -2.7e-06 [-1.0e-05, 4.7e-06] | -5.1e-06 [-1.3e-05, 2.3e-06] | 7.3e-06 [-3.8e-07, 1.5e-05] |
| blaKPC | **-1.5e-05 [-2.9e-05, -8.4e-07]** | -9.6e-06 [-2.4e-05, 5.0e-06] | **-1.7e-05 [-3.2e-05, -2.7e-06]** | -9.3e-06 [-2.4e-05, 5.2e-06] | **-1.6e-05 [-3.1e-05, -1.3e-06]** | -7.4e-06 [-2.3e-05, 8.6e-06] |
| blaOXA | -2.5e-06 [-6.4e-06, 1.5e-06] | -2.6e-06 [-6.7e-06, 1.4e-06] | -2.7e-06 [-6.7e-06, 1.3e-06] | -7.0e-07 [-4.7e-06, 3.3e-06] | -1.8e-06 [-5.9e-06, 2.2e-06] | 1.5e-06 [-2.7e-06, 5.7e-06] |
| blaSHV | 1.9e-06 [3.3e-07, 3.6e-06] | **2.5e-06 [9.6e-07, 4.1e-06]** | **2.6e-06 [9.9e-07, 4.1e-06]** | -3.8e-07 [-2.1e-06, 1.3e-06] | -6.1e-07 [-2.3e-06, 1.1e-06] | 2.6e-07 [-1.6e-06, 2.1e-06] |
| blaTEM | -8.0e-05 [-1.7e-04, 8.5e-06] | -3.4e-05 [-1.2e-04, 5.5e-05] | -7.5e-05 [-1.6e-04, 1.4e-05] | -8.7e-05 [-1.8e-04, 2.0e-06] | **-1.2e-04 [-2.1e-04, -3.1e-05]** | 4.7e-05 [-5.4e-05, 1.5e-04] |
| blaVIM | -6.8e-07 [-2.7e-06, 1.4e-06] | -1.6e-07 [-2.2e-06, 1.9e-06] | 4.4e-07 [-1.6e-06, 2.5e-06] | 1.0e-06 [-1.0e-06, 3.1e-06] | -1.1e-07 [-2.2e-06, 2.0e-06] | -6.1e-07 [-2.7e-06, 1.4e-06] |

| Gene | min °C | max °C | water | Precipitation D2 | Precipitation D1 | Precipitation D0 |
|---|---|---|---|---|---|---|
| ermB | -2.2e-04 [-6.0e-04, 1.5e-04] | -2.1e-04 [-5.9e-04, 1.6e-04] | -1.8e-04 [-5.6e-04, 2.0e-04] | -1.4e-04 [-5.2e-04, 2.4e-04] | -1.8e-04 [-5.7e-04, 2.1e-04] | -1.6e-05 [-4.4e-04, 4.1e-04] |
| intI1 | -2.6e-06 [-2.8e-05, 2.3e-05] | -4.7e-07 [-2.7e-05, 2.6e-05] | -5.4e-06 [-3.2e-05, 2.1e-05] | -7.1e-06 [-3.3e-05, 1.9e-05] | -1.8e-05 [-4.5e-05, 8.9e-06] | 3.4e-06 [-2.6e-05, 3.2e-05] |
| mecA | -2.5e-06 [-5.4e-06, 4.1e-07] | -2.3e-06 [-5.2e-06, 6.8e-07] | -1.7e-06 [-4.8e-06, 1.3e-06] | -1.9e-06 [-4.9e-06, 1.0e-06] | -1.3e-06 [-4.3e-06, 1.7e-06] | -2.3e-06 [-5.6e-06, 9.5e-07] |
| qnrS | **-3.7e-04 [-7.4e-04, -8.5e-06]** | **-4.1e-04 [-7.7e-04, -4.2e-05]** | **-5.3e-04 [-8.9e-04, -1.7e-04]** | -2.0e-04 [-5.7e-04, 1.8e-04] | -3.7e-04 [-7.5e-04, 6.0e-06] | **5.7e-04 [1.7e-04, 9.7e-04]** |
| sul1 | 4.1e-04 [-1.3e-04, 9.5e-04] | 5.0e-04 [-3.7e-05, 1.0e-03] | 5.2e-04 [-1.2e-05, 1.1e-03] | -7.9e-05 [-6.3e-04, 4.7e-04] | -6.5e-05 [-6.3e-04, 5.0e-04] | -4.3e-04 [-1.0e-03, 1.5e-04] |
| tetM | -1.1e-04 [-2.4e-04, 1.5e-05] | -5.1e-05 [-1.8e-04, 8.0e-05] | -1.2e-04 [-2.5e-04, 8.2e-06] | -9.9e-05 [-2.3e-04, 3.3e-05] | -1.3e-04 [-2.6e-04, 5.2e-06] | 7.7e-05 [-6.9e-05, 2.2e-04] |

**Supplementary Table S9. Univariate results for each gene of clinical interest.**

Estimated coefficients from univariate analyses, using the mixed-effect model.
Coefficients associated with a p-value < 5% are highlighted in bold.

| Gene | Δ Macrolides | Δ Sulfaguanidine | Δ Biocides | Δ Heavy Metals | Δ Anti-inflammatory | Δ Microbiome | Maximum Temperature |
|---|---|---|---|---|---|---|---|
| aac(6')-l B | -1.4e-04 [-8.0e-04, 5.1e-04] | **9.0e-04 [2.4e-04, 1.6e-03]** | -6.3e-04 [-1.3e-03, 6.3e-05] | 3.6e-04 [-3.5e-04, 1.1e-03] | **-8.0e-04 [-1.4e-03, -1.9e-04]** | **-1.2e-03 [-1.8e-03, -5.3e-04]** | -2.3e-04 [-8.4e-04, 3.8e-04] |
| aph(3')-III | 2.0e-05 [-1.4e-05, 5.5e-05] | 1.9e-05 [-1.6e-05, 5.5e-05] | 2.6e-05 [-1.0e-05, 6.3e-05] | 4.3e-06 [-3.3e-05, 4.2e-05] | -3.2e-05 [-6.6e-05, 1.2e-06] | **-7.2e-05 [-1.0e-04, -4.0e-05]** | -7.9e-06 [-4.1e-05, 2.6e-05] |
| blaCTX-M | -2.9e-06 [-1.1e-05, 5.0e-06] | 2.8e-07 [-7.6e-06, 8.1e-06] | -8.4e-07 [-9.3e-06, 7.6e-06] | **7.9e-06 [3.2e-07, 1.6e-05]** | **-1.0e-05 [-1.8e-05, -3.2e-06]** | 1.3e-06 [-6.6e-06, 9.2e-06] | **-1.2e-05 [-1.9e-05, -5.7e-06]** |
| blaKPC | -2.4e-06 [-1.9e-05, 1.4e-05] | 5.4e-06 [-1.1e-05, 2.2e-05] | 5.6e-06 [-1.2e-05, 2.3e-05] | 5.2e-06 [-1.2e-05, 2.3e-05] | -1.4e-05 [-2.9e-05, 2.3e-06] | **-1.7e-05 [-3.3e-05, -7.8e-07]** | -9.6e-06 [-2.4e-05, 5.0e-06] |
| blaOXA | **-4.2e-06 [-8.3e-06, -3.0e-08]** | 1.5e-06 [-2.8e-06, 5.8e-06] | 2.4e-06 [-1.9e-06, 6.7e-06] | 1.5e-06 [-2.8e-06, 5.8e-06] | **4.4e-06 [3.1e-07, 8.4e-06]** | 1.4e-06 [-2.9e-06, 5.6e-06] | -2.6e-06 [-6.7e-06, 1.4e-06] |
| blaSHV | 1.4e-06 [-4.4e-07, 3.2e-06] | 3.4e-07 [-1.5e-06, 2.2e-06] | -5.5e-07 [-2.5e-06, 1.4e-06] | 6.6e-07 [-1.4e-06, 2.7e-06] | -1.9e-07 [-2.0e-06, 1.6e-06] | **-4.2e-06 [-5.8e-06, -2.7e-06]** | **2.5e-06 [9.6e-07, 4.1e-06]** |
| blaTEM | 4.9e-05 [-4.8e-05, 1.5e-04] | 7.3e-05 [-2.8e-05, 1.7e-04] | -5.8e-06 [-1.1e-04, 9.8e-05] | 5.2e-05 [-5.5e-05, 1.6e-04] | -6.4e-05 [-1.6e-04, 2.8e-05] | **-2.3e-04 [-3.2e-04, -1.4e-04]** | -3.4e-05 [-1.2e-04, 5.5e-05] |
| blaVIM | 3.4e-07 [-2.0e-06, 2.7e-06] | 1.5e-06 [-8.8e-07, 3.8e-06] | 7.3e-07 [-1.6e-06, 3.1e-06] | 1.4e-06 [-9.7e-07, 3.7e-06] | -1.9e-06 [-4.2e-06, 4.3e-07] | -9.9e-07 [-3.3e-06, 1.4e-06] | -1.6e-07 [-2.2e-06, 1.9e-06] |
| ermB | **7.9e-04 [4.2e-04, 1.2e-03]** | 1.0e-05 [-4.1e-04, 4.3e-04] | **5.4e-04 [1.2e-04, 9.6e-04]** | 3.3e-04 [-1.1e-04, 7.7e-04] | -3.0e-04 [-6.8e-04, 8.5e-05] | -9.8e-05 [-5.3e-04, 3.3e-04] | -2.1e-04 [-5.9e-04, 1.6e-04] |
| intI1 | **4.6e-05 [2.0e-05, 7.3e-05]** | 7.4e-06 [-2.2e-05, 3.7e-05] | 1.8e-05 [-1.5e-05, 5.2e-05] | 1.8e-05 [-1.2e-05, 4.7e-05] | **-3.1e-05 [-5.7e-05, -4.7e-06]** | **-4.4e-05 [-7.1e-05, -1.6e-05]** | -4.7e-07 [-2.7e-05, 2.6e-05] |
| mecA | -2.2e-06 [-5.4e-06, 1.1e-06] | **3.7e-06 [4.9e-07, 7.0e-06]** | -4.5e-07 [-3.8e-06, 2.9e-06] | 1.5e-06 [-1.9e-06, 4.9e-06] | 1.2e-06 [-1.9e-06, 4.4e-06] | -1.7e-06 [-5.2e-06, 1.9e-06] | -2.3e-06 [-5.2e-06, 6.8e-07] |
| qnrS | 2.3e-04 [-1.7e-04, 6.3e-04] | 3.9e-04 [-2.0e-05, 8.0e-04] | -1.5e-04 [-5.8e-04, 2.7e-04] | 5.1e-05 [-3.9e-04, 4.9e-04] | -3.7e-04 [-7.4e-04, 1.1e-05] | **-6.1e-04 [-1.0e-03, -2.1e-04]** | **-4.1e-04 [-7.7e-04, -4.2e-05]** |
| sul1 | -8.2e-05 [-6.6e-04, 4.9e-04] | **7.2e-04 [1.4e-04, 1.3e-03]** | 2.3e-04 [-3.5e-04, 8.1e-04] | 2.8e-04 [-3.1e-04, 8.6e-04] | -2.6e-04 [-8.2e-04, 3.0e-04] | **-1.6e-03 [-2.0e-03, -1.1e-03]** | 5.0e-04 [-3.7e-05, 1.0e-03] |

| Gene | Δ Macrolides | Δ Sulfaguanidine | Δ Biocides | Δ Heavy Metals | Δ Anti-inflammatory | Δ Microbiome | Maximum Temperature |
|---|---|---|---|---|---|---|---|
| tetM | **1.7e-04 [3.3e-05, 3.1e-04]** | 8.2e-05 [-6.6e-05, 2.3e-04] | 9.6e-05 [-5.5e-05, 2.5e-04] | 3.7e-05 [-1.2e-04, 1.9e-04] | **-1.7e-04 [-3.0e-04, -3.8e-05]** | **-2.3e-04 [-3.7e-04, -9.2e-05]** | -5.1e-05 [-1.8e-04, 8.0e-05] |

**Supplementary Table S10. Multivariate results for each gene of clinical interest.**

Estimated coefficients from univariate analyses, using the mixed-effect model. Coefficients associated with a p-value < 5% are highlighted in bold.

| Gene | Δ Macrolides | Δ Sulfaguanidine | Δ Biocides | Δ Heavy Metals | Δ Anti-inflammatory | Δ Microbiome (log) | Maximum Temperature |
|---|---|---|---|---|---|---|---|
| aac(6')-l B | -1.1e-04<br>[-7.2e-04, 5.0e-04] | **6.6e-04<br>[2.9e-05, 1.3e-03]** | -4.5e-04<br>[-1.1e-03, 2.4e-04] | 2.6e-04<br>[-3.8e-04, 9.0e-04] | **-6.3e-04<br>[-1.2e-03, -4.3e-05]** | **-1.2e-03<br>[-1.9e-03, -5.9e-04]** | -4.1e-04<br>[-9.8e-04, 1.6e-04] |
| aph(3')-III | 1.2e-05<br>[-2.2e-05, 4.7e-05] | 2.0e-05<br>[-1.3e-05, 5.3e-05] | 2.4e-05<br>[-1.3e-05, 6.2e-05] | 1.1e-05<br>[-2.4e-05, 4.5e-05] | **-3.4e-05<br>[-6.7e-05, -2.0e-07]** | **-6.7e-05<br>[-1.0e-04, -3.2e-05]** | -6.5e-06<br>[-3.9e-05, 2.6e-05] |
| blaCTX-M | 2.3e-08<br>[-8.0e-06, 8.0e-06] | 3.1e-07<br>[-7.6e-06, 8.2e-06] | -6.9e-07<br>[-9.9e-06, 8.5e-06] | 6.8e-06<br>[-9.7e-07, 1.5e-05] | -6.4e-06<br>[-1.4e-05, 1.1e-06] | -2.4e-06<br>[-1.1e-05, 5.7e-06] | **-1.2e-05<br>[-1.9e-05, -4.4e-06]** |
| blaKPC | -3.0e-06<br>[-1.9e-05, 1.3e-05] | 8.6e-06<br>[-9.0e-06, 2.6e-05] | 6.5e-06<br>[-1.3e-05, 2.6e-05] | 4.6e-06<br>[-1.3e-05, 2.2e-05] | -1.2e-05<br>[-2.8e-05, 4.1e-06] | **-1.9e-05<br>[-3.6e-05, -2.1e-06]** | -1.1e-05<br>[-2.7e-05, 4.4e-06] |
| blaOXA | **-5.3e-06<br>[-9.5e-06, -1.1e-06]** | 1.9e-06<br>[-2.3e-06, 6.2e-06] | 2.6e-06<br>[-2.1e-06, 7.3e-06] | 2.1e-06<br>[-2.0e-06, 6.3e-06] | **4.9e-06<br>[7.9e-07, 9.1e-06]** | 2.7e-07<br>[-4.0e-06, 4.6e-06] | -1.8e-06<br>[-6.0e-06, 2.4e-06] |
| blaSHV | 6.0e-07<br>[-1.4e-06, 2.6e-06] | -2.5e-07<br>[-2.3e-06, 1.8e-06] | -5.9e-07<br>[-2.9e-06, 1.7e-06] | 1.1e-06<br>[-9.4e-07, 3.2e-06] | -2.9e-08<br>[-1.9e-06, 1.8e-06] | **-3.7e-06<br>[-5.7e-06, -1.7e-06]** | 1.9e-06<br>[-5.7e-08, 3.8e-06] |
| blaTEM | 3.1e-05<br>[-5.9e-05, 1.2e-04] | 4.2e-05<br>[-5.2e-05, 1.4e-04] | -1.8e-05<br>[-1.2e-04, 8.4e-05] | 4.9e-05<br>[-4.7e-05, 1.4e-04] | -5.0e-05<br>[-1.4e-04, 3.5e-05] | **-2.4e-04<br>[-3.3e-04, -1.4e-04]** | -6.7e-05<br>[-1.5e-04, 1.6e-05] |
| blaVIM | 1.7e-07<br>[-2.4e-06, 2.7e-06] | 6.6e-07<br>[-1.8e-06, 3.2e-06] | 1.8e-06<br>[-9.9e-07, 4.6e-06] | 1.9e-06<br>[-6.0e-07, 4.4e-06] | -2.5e-06<br>[-5.0e-06, 1.2e-07] | -1.1e-06<br>[-3.7e-06, 1.5e-06] | -9.0e-08<br>[-2.6e-06, 2.4e-06] |
| ermB | **7.8e-04<br>[4.1e-04, 1.2e-03]** | 8.5e-05<br>[-3.1e-04, 4.8e-04] | **4.6e-04<br>[3.0e-05, 8.8e-04]** | 2.4e-04<br>[-1.6e-04, 6.4e-04] | **-4.7e-04<br>[-8.2e-04, -1.1e-04]** | 3.6e-05<br>[-3.5e-04, 4.2e-04] | -1.6e-04<br>[-5.0e-04, 1.9e-04] |
| intI1 | **3.9e-05<br>[1.3e-05, 6.5e-05]** | 6.7e-06<br>[-2.3e-05, 3.6e-05] | 1.5e-05<br>[-2.0e-05, 5.0e-05] | 1.6e-05<br>[-1.0e-05, 4.3e-05] | **-3.5e-05<br>[-6.0e-05, -1.1e-05]** | **-3.7e-05<br>[-6.3e-05, -1.1e-05]** | -3.5e-06<br>[-2.8e-05, 2.1e-05] |
| mecA | -2.2e-06<br>[-5.5e-06, 1.2e-06] | **3.7e-06<br>[2.4e-07, 7.1e-06]** | 2.6e-07<br>[-3.3e-06, 3.8e-06] | 1.1e-06<br>[-2.2e-06, 4.5e-06] | 2.2e-06<br>[-1.1e-06, 5.4e-06] | -1.1e-06<br>[-4.8e-06, 2.6e-06] | -2.7e-06<br>[-5.8e-06, 3.4e-07] |

| Gene | Δ Macrolides | Δ Sulfaguanidine | Δ Biocides | Δ Heavy Metals | Δ Anti-inflammatory | Δ Microbiome (log) | Maximum Temperature |
|---|---|---|---|---|---|---|---|
| qnrS | 2.5e-04 [-1.2e-04, 6.3e-04] | 3.2e-04 [-6.8e-05, 7.2e-04] | -2.4e-04 [-6.7e-04, 1.9e-04] | 4.7e-05 [-3.5e-04, 4.5e-04] | -3.0e-04 [-6.6e-04, 6.3e-05] | **-5.7e-04 [-9.6e-04, -1.9e-04]** | **-6.0e-04 [-9.6e-04, -2.5e-04]** |
| sul1 | **-7.8e-04 [-1.2e-03, -3.4e-04]** | **6.7e-04 [2.2e-04, 1.1e-03]** | **4.7e-04 [1.4e-05, 9.2e-04]** | 3.2e-04 [-1.2e-04, 7.6e-04] | -3.2e-04 [-7.4e-04, 1.1e-04] | **-1.6e-03 [-2.1e-03, -1.2e-03]** | **5.8e-04 [1.7e-04, 9.9e-04]** |
| tetM | **1.6e-04 [2.5e-05, 2.9e-04]** | 9.6e-05 [-4.4e-05, 2.3e-04] | 8.7e-05 [-6.3e-05, 2.4e-04] | 2.3e-05 [-1.2e-04, 1.7e-04] | **-1.9e-04 [-3.2e-04, -6.5e-05]** | **-1.9e-04 [-3.3e-04, -5.9e-05]** | -1.1e-04 [-2.3e-04, 1.7e-05] |

**Supplementary Table S11. Multivariable versus null model AIC and spatial random effects variance for each gene of clinical interest.**

*ΔAIC = AIC_null - AIC_full

Bold values indicate models for which the full model has a lower AIC than the null model

| Gene | AIC Null | AIC Full | ΔAIC* | Var. Sampling Point (Null) | Var. Sampling Point (Full) |
|---|---|---|---|---|---|
| acc(6')-1 B | -712 | - 729 | **16.7** | 4.02e-06 | 2.77e-06 |
| aph (3')-III | -1129 | -1140 | **10.6** | 3.16e-09 | 0 |
| blaCTX-M | - 1189 | - 1195 | **6.1** | 2.03e-10 | 1.94e-10 |
| blaKPC | - 1076 | - 1074 | -2.4 | 2.65e-09 | 2.17e-09 |
| blaOXA | - 1366 | - 1366 | **0.5** | 3.20e-11 | 2.28e-11 |
| blaSHV | -1423 | -1442 | **19.2** | 3.11e-11 | 2.51e-11 |
| blaTEM | -1029 | -1046 | **17.1** | 3.44e-07 | 2.71e-07 |
| blaVIM | -1233 | -1229 | -3.9 | 0 | 4.35e-29 |
| ermB | -793 | -811 | **17.1** | 4.59e-06 | 3.61e-06 |
| intI1 | -1124 | -1139 | **15.4** | 2.20e-09 | 1.56e-08 |
| mecA | -1142 | -1141 | -1.6 | 3.80e-11 | 1.92e-11 |
| qnrS | -801 | -813 | **12.1** | 1.85e-06 | 1.17e-06 |
| sul1 | - 680 | -726 | **46.3** | 1.46e-06 | 5.33e-07 |
| tetM | -968 | -983 | **14.3** | 4.95e-07 | 2.93e-07 |

**Supplementary Figure S13. Violin plots of spatial random effects for each gene of clinical interest.**

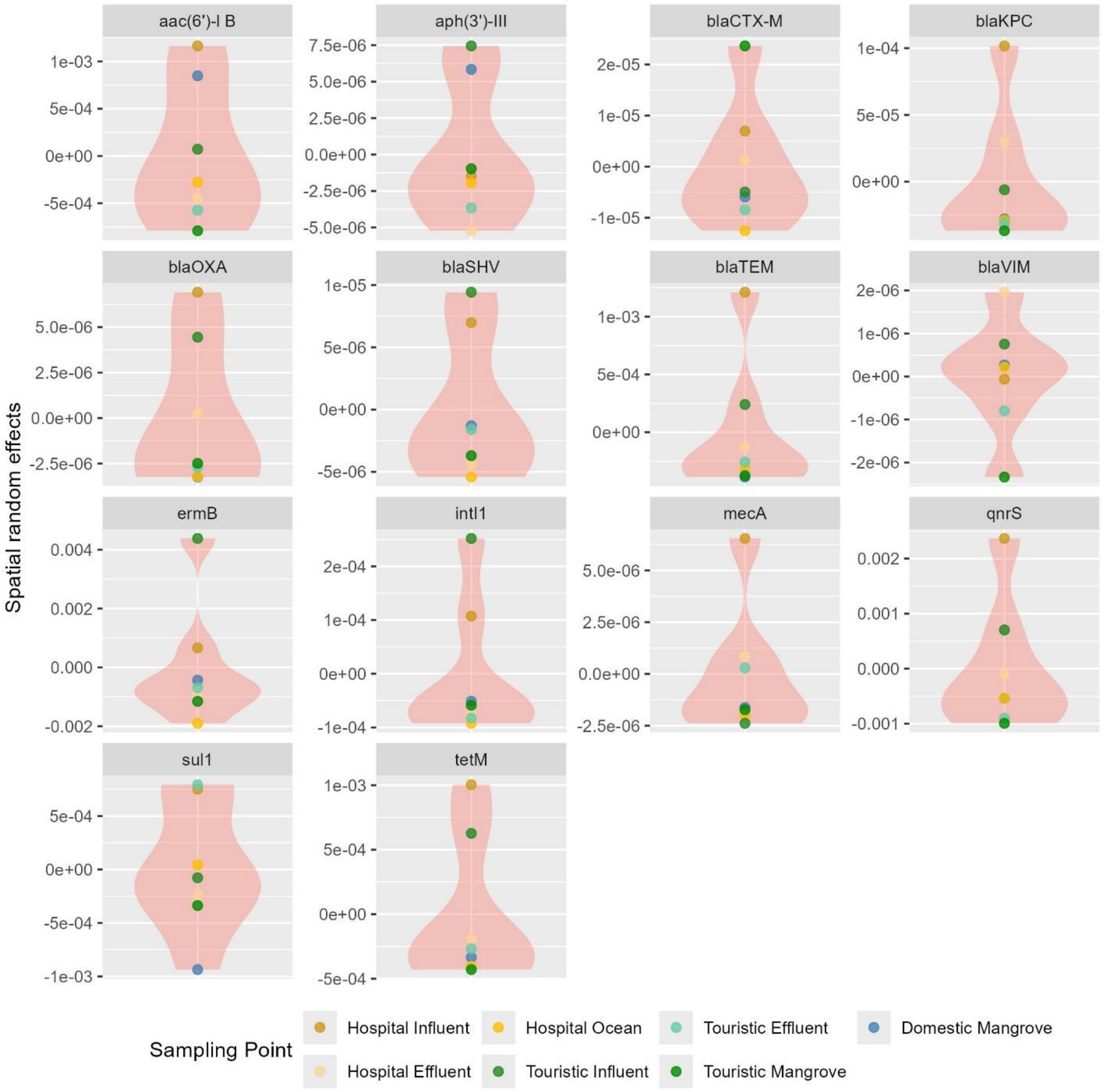

**Supplementary Table S12. Multivariable versus null model AIC and spatial random effects variance for the ESKAPEE-associated genera.**

*ΔAIC = AIC_null - AIC_full

Bold values indicate models for which the full model has a lower AIC than the null model

| **Genus** | ΔAIC * | Var. Sampling Point (Null) | Var. Sampling Point (Full) |
|---|---|---|---|
| *Enterococcus* | **1.03** | 9.53e-6 (98%) | 0 (0%) |
| *Staphylococcus* | **8.20** | 1.15e-7 (42%) | 0 (0%) |
| *Klebsiella* | -8.60 | 8.35e-7 (68%) | 0 (0%) |
| *Acinetobacter* | -3.61 | 3.75e-4 (94%) | 1.07e-4 (81%) |
| *Pseudomonas* | -1.12 | 0 (0%) | 0 (0%) |
| *Enterobacter* | -5.75 | 4.09e-8 (57%) | 1.24e-8 (29%) |
| *Escherichia* | -9.10 | 1.42e-10 (27%) | 1.25e-10 (26%) |

**Supplementary Figure S14. Violin plots of spatial random effects for the ESKAPEE-associated genera.**

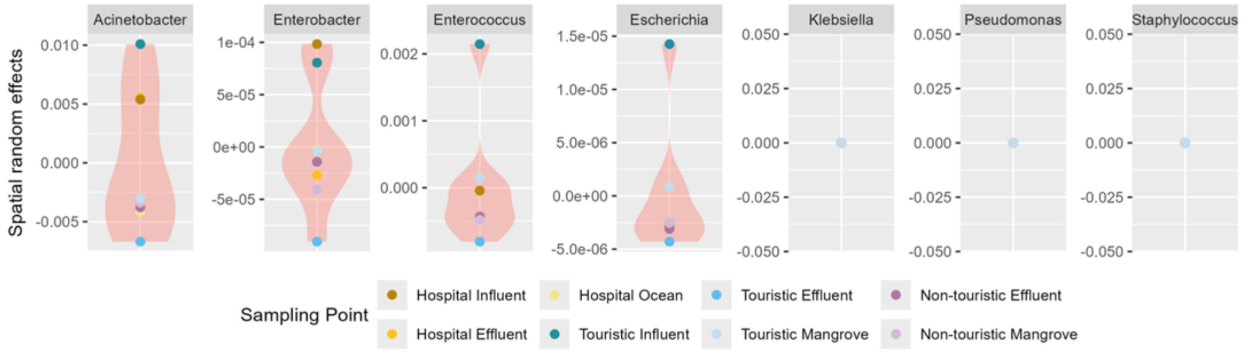

**Supplementary Figure S15. Estimated effects of covariates on the abundance of ESKAPEE-associated genera from the mixed-effects models.** Points show regression coefficients from 7 mixed-effects models (one per ESKAPEE genera), with horizontal bars representing 95% credible intervals. The dashed vertical line indicates no effect ($\beta = 0$). Blue and red denote negative and positive associations, respectively. Statistical significance is indicated by asterisks (*p < 0.05; **p < 0.01; ***p < 0.001. *ΔMicrobiome* corresponds to the Bray–Curtis dissimilarity between upstream and downstream sites. Other covariates are expressed as downstream minus upstream values.

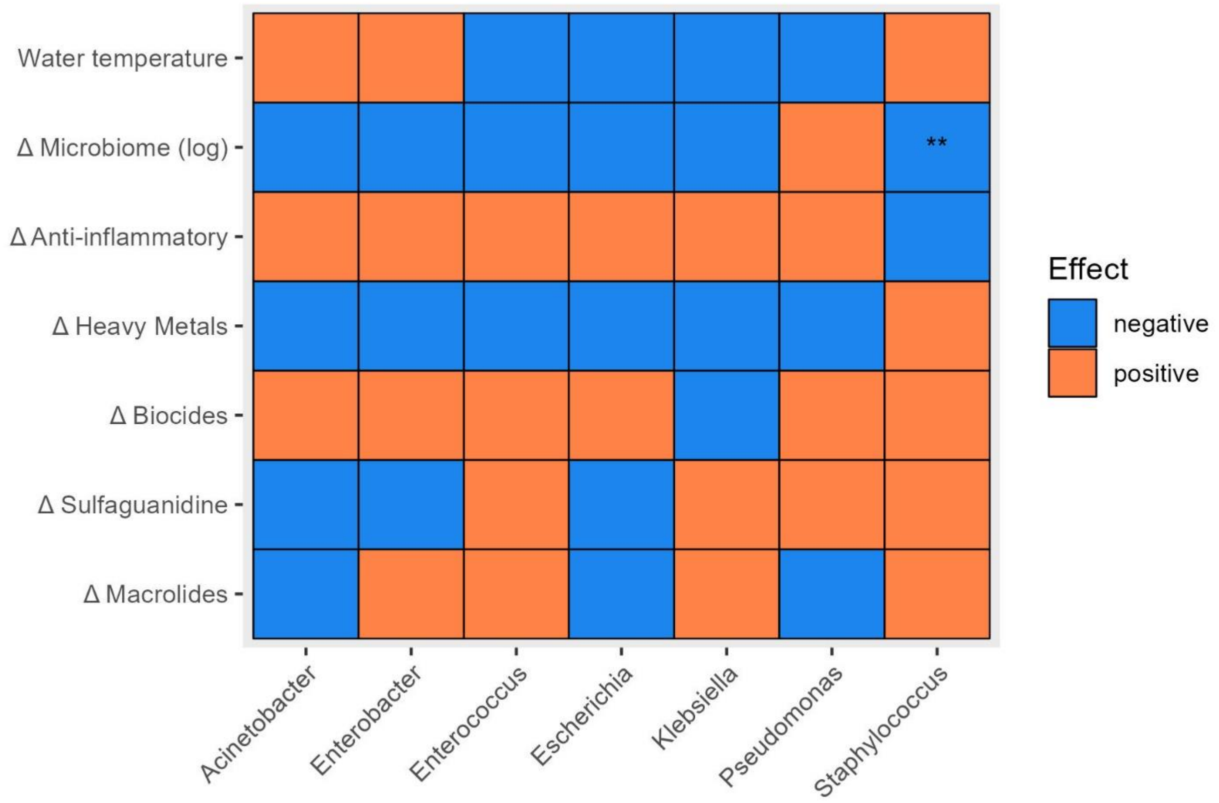

### Sensitivity analysis

*Description of the results*

Only three genes had an AIC in the full model higher than in the null model *( aph(3')-III, ermB, sul1).*

**Supplementary Table S13. Multivariable versus null model AIC and spatial random effects variance for the sensitivity analysis.**

*ΔAIC = AIC_null - AIC_full

Bold values indicate models for which the full model has a lower AIC than the null model

| Gene | AIC Null | AIC Full | ΔAIC* |
|---|---|---|---|
| aac(6')-I B | -540.0 | -531.0 | -9.78 |
| aph(3')-III | -971.0 | -977.0 | **5.69** |

| | | | |
|---|---|---|---|
| blaCTX-M | -834.0 | -829.0 | -5.23 |
| blaKPC | -811.0 | -800.0 | -10.3 |
| blaOXA | -1133.0 | -1125.0 | -7.51 |
| blaSHV | -1255.0 | -1249.0 | -4.48 |
| blaTEM | -901.0 | -892.0 | -8.39 |
| blaVIM | -582.0 | -569.5 | -13.1 |
| ermB | -722.0 | -722.0 | **0.0296** |
| intI1 | -1007.0 | -1002.0 | -4.74 |
| mecA | -1127.0 | -1126.0 | 1.32 |
| qnrS | -638.0 | -630.0 | -8.24 |
| sul1 | -653.0 | -656.0 | **3.55** |
| tetM | -933.0 | -926.0 | -7.14 |

**Supplementary Figure S16. Estimated effects of covariates on the difference in abundance of ARGs from the mixed-effects models.** Points show regression coefficients from the mixed-effects models, with horizontal bars representing 95% credible intervals. The dashed vertical line indicates no effect ($\boldsymbol{\beta}$ = 0). Blue and red denote negative and positive associations, respectively. Statistical significance is indicated by asterisks (*p < 0.05; **p < 0.01; ***p < 0.001. $\boldsymbol{\Delta}$*Microbiome* corresponds to the Bray–Curtis dissimilarity between upstream and downstream sites. Other covariates are expressed as downstream minus upstream values.

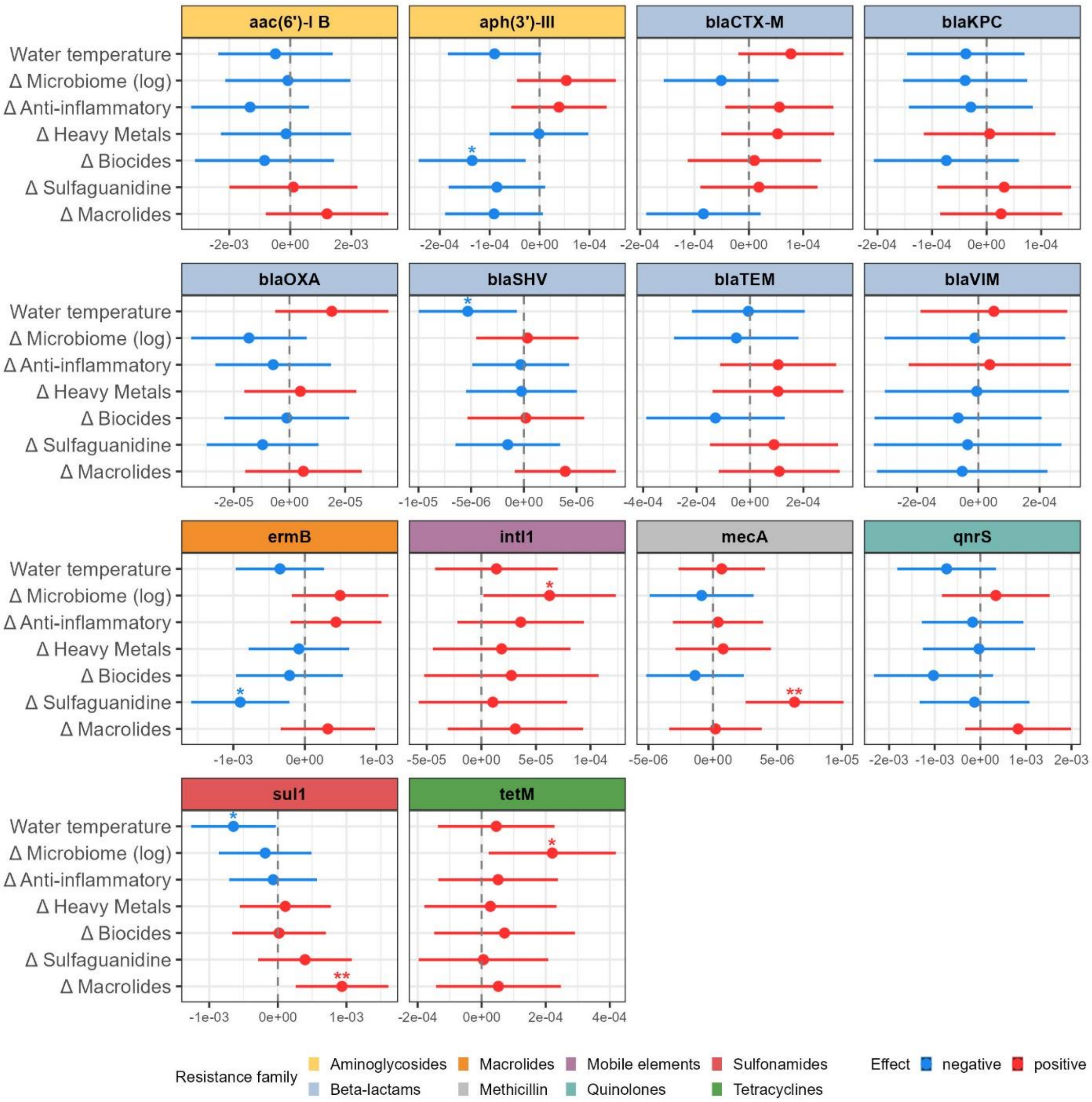
aac(6')-I B
aph(3')-III
blaCTX-M
blaKPC
blaOXA
blaSHV
blaTEM
blaVIM
ermB
intI1
mecA
qnrS
sul1
tetM
Water temperature
Δ Microbiome (log)
Δ Anti-inflammatory
Δ Heavy Metals
Δ Biocides
Δ Sulfaguanidine
Δ Macrolides
-2e-03 0e+00 2e-03
-2e-04 -1e-04 0e+00 1e-04
-2e-04 -1e-04 0e+00 1e-04
-2e-04 -1e-04 0e+00 1e-04
-2e-05 0e+00 2e-05
-1e-05 -5e-06 0e+00 5e-06
-4e-04 -2e-04 0e+00 2e-04
-2e-04 0e+00 2e-04
-1e-03 0e+00 1e-03
-5e-05 0e+00 5e-05 1e-04
-5e-06 0e+00 5e-06 1e-05
-2e-03 -1e-03 0e+00 1e-03 2e-03
-1e-03 0e+00 1e-03
-2e-04 0e+00 2e-04 4e-04
Resistance family
Aminoglycosides
Beta-lactams
Macrolides
Methicillin
Mobile elements
Quinolones
Sulfonamides
Tetracyclines
Effect
negative
positive
